# Characterization of Primed Adaptation in the *Escherichia coli* type I-E CRISPR-Cas System

**DOI:** 10.1101/2020.02.10.942821

**Authors:** Anne M. Stringer, Lauren A. Cooper, Sujatha Kadaba, Shailab Shrestha, Joseph T. Wade

## Abstract

CRISPR-Cas systems are bacterial immune systems that target invading nucleic acid. The hallmark of CRISPR-Cas systems is the CRISPR array, a genetic locus that includes short sequences known as “spacers”, that are derived from invading nucleic acid. Upon exposure to an invading nucleic acid molecule, bacteria/archaea with functional CRISPR-Cas systems can add new spacers to their CRISPR arrays in a process known as “adaptation”. In type I CRISPR-Cas systems, which represent the majority of CRISPR-Cas systems found in nature, adaptation can occur by two mechanisms: naïve and primed. Here, we show that, for the archetypal type I-E CRISPR-Cas system from *Escherichia coli*, primed adaptation occurs at least 1,000 times more efficiently than naïve adaptation. By initiating primed adaptation on the *E. coli* chromosome, we show that spacers can be acquired across distances of >100 kb from the initially targeted site, and we identify multiple factors that influence the efficiency with which sequences are acquired as new spacers. Thus, our data provide insight into the mechanism of primed adaptation.

[*This paper has been peer reviewed, with Ailong Ke (Cornell University) serving as the editor. Reviews and point-by-point response, and a marked-up version of the edited manuscript are provided as supplementary files*.]

## INTRODUCTION

CRISPR-Cas systems are adaptive immune systems that are widespread throughout the bacterial and archaeal kingdoms and protect cells from invading nucleic acid molecules. Although the mechanistic details vary widely across different CRISPR-Cas systems, the basic mechanism is shared across all systems (Wright et al., 2016). A short RNA, known as a CRISPR RNA (crRNA) associates with a surveillance CRISPR-associated (Cas) protein, or complex of Cas proteins. This protein-RNA complex binds to the invading nucleic acid molecule, at least in part due to base-pairing between the crRNA and complementary sequence in the invading nucleic acid (the “protospacer”). This leads to cleavage of the invading nucleic acid either by the surveillance protein-RNA complex, or by an additional protein that is recruited to this complex. The most abundant type of CRISPR-Cas system in nature is the type I system (Koonin et al., 2017); the archetypal type I system is the type I-E system of *Escherichia coli*, which employs a large complex of five Cas proteins (some proteins present in multiple copies), “Cascade”, as the surveillance complex, and recruits a separate nuclease protein, Cas3 (Brouns et al., 2008).

A hallmark of all CRISPR-Cas systems is the CRISPR array, a genetic locus consisting of multiple units of short (∼20-40 bp) “repeat” and “spacer” sequences (Wright et al., 2016). Within a given CRISPR array, repeat sequences are identical, and each repeat is separated from the next by a spacer. Spacer sequences are variable, and are derived from invading nucleic acid molecules. The CRISPR array is typically transcribed as a single, long RNA, which is then processed to yield individual crRNAs. Each crRNA contains a single spacer flanked by partial repeats, and the spacer sequence provides the base-pairing complementarity to a protospacer in the nucleic acid target. For CRISPR-Cas systems targeting DNA, an additional sequence in the DNA target is typically required for CRISPR-Cas immunity. This short sequence (typically 2-4 bp), known as the protospacer-adjacent motif (PAM), is located close to the protospacer, and serves as a binding site for one of the Cas proteins in the surveillance complex (Leenay and Beisel, 2017). There is often some flexibility in the PAM sequences, with multiple sequences being sufficient to recruit Cas proteins. However, there are typically one or a few “optimal” PAM sequences that have the highest affinity for the cognate Cas protein.

A key feature of all CRISPR-Cas systems is their ability to acquire new spacers, in a process known as adaptation (Jackson et al., 2017; Sternberg et al., 2016). During adaptation, one or more spacer+repeat units are added to the CRISPR array. Spacer+repeat units are typically added at one end of the array, due to the presence of a flanking sequence known as the leader (Yosef et al., 2012). Adaptation requires the Cas1 and Cas2 proteins, which, in *E. coli*, form a hexameric complex (Cas14-Cas22; referred to here as “Cas1-Cas2”) that binds dsDNA fragments containing a PAM (Nuñez et al., 2014, 2015a; Wang et al., 2015; Yosef et al., 2012). Cas1-Cas2 catalyzes nucleophilic attack of the CRISPR array by the 3’-OH groups of the bound DNA fragment (Nuñez et al., 2015b). Recent studies suggest that in *Escherichia coli*, Cas1 binding to the PAM protects the PAM-proximal end of the bound DNA from exonucleolytic digestion, whereas the PAM-distal end is not protected. The resulting difference in 3’ overhang length dictates the order of nucleophilic attack, and thus ensures that the PAM-proximal side is integrated closest to the leader (Drabavicius et al., 2018; Kim et al., 2020; Ramachandran et al., 2020).

For type I CRISPR-Cas systems, there are two mechanisms of adaptation: naïve and primed. Naïve adaptation occurs when an organism is invaded by a nucleic acid sequence that is not already matched by an existing spacer. Naïve adaptation typically requires only the Cas1 and Cas2 proteins, and has been proposed to occur by Cas1-Cas2 acquisition of DNA fragments generated by resection of double-strand breaks (Levy et al., 2015). By contrast, primed adaptation requires invasion of the cell by a nucleic acid sequence for which there is already a partially or completely matching spacer. The mechanism of primed adaptation is not well understood, but requires all Cas proteins (Datsenko et al., 2012). Targeting of a protospacer by Cascade with a partially or fully complementary spacer, and an optimal or sub-optimal (but not inactive) PAM, leads to recruitment of Cas3, which is both a nuclease and a helicase (Mulepati and Bailey, 2013; Sinkunas et al., 2011, 2013). Primed adaptation likely involves translocation of Cas3 away from the protospacer (Dillard et al., 2018; Redding et al., 2015) either by Cas3 moving along the DNA or by Cas3 reeling in the DNA while remaining tethered to the protospacer-bound Cascade (Dillard et al., 2018; Loeff et al., 2018; Redding et al., 2015), with Cas3 generating the substrates for Cas1-Cas2 to integrate new spacers into the CRISPR array (Dillard et al., 2018; Künne et al., 2016; Semenova et al., 2016). However, the mechanism by which Cas3 and/or other proteins generate substrates for Cas1-Cas2 is poorly understood (Musharova et al., 2017). A recent study examined the nature of DNA fragments bound by Cas1-Cas2 in *E. coli* undergoing primed adaptation. Surprisingly, the DNA fragments did not have the ends expected based on biochemical studies of adaptation (Kim et al., 2020; Ramachandran et al., 2020), but rather had a blunt PAM-distal end and a 4 nt 3’ overhang at the PAM-proximal end, with the 5’ nucleotide at the PAM-proximal end being the last nucleotide of the PAM. These data suggest that the processing of fragments bound by Cas1-Cas2 may be different during naïve and primed adaptation.

For the purposes of this study, we refer to the source of a spacer as the “pre-spacer” location, and we refer to the site of Cascade binding as the protospacer. The sequence context of spacers acquired during primed adaptation has a large impact on the efficiency with which a given sequence is acquired (Datsenko et al., 2012; Fineran et al., 2014; Li et al., 2014; Musharova et al., 2018, 2018; Rao et al., 2017; Richter et al., 2014; Savitskaya et al., 2013; Shmakov et al., 2014; Staals et al., 2016; Strotskaya et al., 2017; Swarts et al., 2012). Previous studies have shown that pre-spacers are located predominantly upstream of the protospacer, consistent with the unidirectional helicase activity of Cas3. For type I-F systems, pre-spacers are equally abundant on both the target and non-target strands (Richter et al., 2014; Staals et al., 2016), whereas for type I-B, I-C, and I-E systems, pre-spacers are located predominantly on the non-target strand (Datsenko et al., 2012; Fineran et al., 2014; Garrett et al., 2020; Li et al., 2014; Rao et al., 2017; Savitskaya et al., 2013; Shmakov et al., 2014; Stachler et al., 2020; Strotskaya et al., 2017; Swarts et al., 2012). For type I-F systems, pre-spacers are located over a ∼5 kb region upstream of the protospacer (Staals et al., 2016), whereas for type I-E systems, acquisition has been reported over distances of up to ∼10 kb (Strotskaya et al., 2017), although most studies involved plasmids of <10 kbp for which the distance of primed adaptation cannot be measured since it likely exceeds the size of the plasmid (Datsenko et al., 2012; Fineran et al., 2014; Savitskaya et al., 2013; Shmakov et al., 2014). For all type I systems tested to date, most pre-spacers are flanked by an optimal PAM sequence (Datsenko et al., 2012; Fineran et al., 2014; Li et al., 2014; Rao et al., 2017; Richter et al., 2014; Savitskaya et al., 2013; Shmakov et al., 2014; Staals et al., 2016; Strotskaya et al., 2017; Swarts et al., 2012); indeed, pre-spacers that are not immediately flanked by an optimal PAM sequence are often the result of (i) “slipping” errors, where an optimal PAM is located one or two bases away from the expected location, suggesting incorrect processing of the DNA by Cas1-Cas2 (Li et al., 2014; Rao et al., 2017; Shmakov et al., 2014; Staals et al., 2016), or (ii) “flipping” errors, where the PAM is on the opposite side and strand of the pre-spacer, suggesting that Cas1-Cas2 has incorrectly inserted the spacer into the CRISPR array in the reverse orientation (Li et al., 2014; Rao et al., 2017; Shmakov et al., 2014; Staals et al., 2016).

Here, we further investigate the process of primed adaptation in the type I-E system of *E. coli*. Previous studies have shown that primed adaptation is considerably more efficient than naïve adaptation in type I-B, I-E and I-F systems (Datsenko et al., 2012; Li et al., 2014; Staals et al., 2016). We quantify this difference for the *E. coli* system, showing that primed adaptation is at least 1,000 times more efficient than naïve adaptation. We observe primed adaptation from the *E. coli* chromosome over distances >100 kb. Using the rich source of pre-spacers inferred from these data, we identify 7 features of pre-spacers that determine the efficiency with which they are acquired, supporting and extending previous observations: (i) the DNA strand of the pre-spacer relative to the protospacer (i.e. target vs non-target strand), (ii) the position of the pre-spacer upstream or downstream of the protospacer, (iii) the presence and position of an AAG PAM immediately adjacent to the pre-spacer, (iv) distance of the pre-spacer from the protospacer over a ∼100 kb region, (v) the presence of an AAG within the pre-spacer, (vi) whether or not the pre-spacer is acquired from within ∼200 bp of the protospacer, (vii) the sequence of positions 30-33 of the pre-spacer. Based on these factors and on prior work from other groups, we propose a model of primed adaptation in which a Cas3-containing complex travels large distances along the DNA from the protospacer, cutting the DNA at or near every AAG, contributing to the generation of substrates for Cas1-Cas2.

## RESULTS AND DISCUSSION

### Primed adaptation is >1,000-fold more efficient than naïve adaptation in *E. coli*

Studies of naïve adaptation in *E. coli* have involved high-level overexpression of Cas1 and Cas2 (Levy et al., 2015; Nuñez et al., 2014; Wang et al., 2015; Yosef et al., 2012, 2013). Hence, it is unclear how the efficiency of naïve adaptation compares to that of primed adaptation. We previously used a highly sensitive, fluorescence-based reporter (Amlinger et al., 2017) to precisely quantify primed adaptation in *E. coli* (Cooper et al., 2018). For protospacers with extensive mismatches in the PAM-proximal “seed” region or the PAM-distal region, we failed to detect adaptation, strongly suggesting that naïve adaptation is extremely inefficient in these cells. To directly compare the efficiencies of primed and naïve adaptation in genetically similar strains, we used a Δ*cas3* strain of *E. coli* in which all other *cas* genes are constitutively expressed from their native locus. This strain also contains a minimal CRISPR array (i.e. one repeat, and a leader sequence) associated with an out-of-frame copy of *yfp*. Expansion of this artificial array by a single repeat/spacer unit puts *yfp* in frame, leading to YFP fluorescence (Figure 1A). While adaptation in the context of this reporter is lower than that of the native CRISPR loci in strains lacking the reporter, perhaps due in part to competition with the native CRISPR-I array, the reporter is nonetheless highly sensitive, allowing detection of very low level adaptation (Cooper et al., 2018). We then introduced a plasmid expressing *cas3* from an arabinose-inducible promoter, or an equivalent empty vector. We also introduced a plasmid expressing a crRNA that targets a protospacer on the same plasmid (“self-targeting plasmid”; stp), or an equivalent empty vector. Thus, we were able to infer the level of adaptation by counting YFP^+^ cells in four different strains: (i) crRNA^-^ *cas3*^-^, (ii) crRNA^+^ *cas3*^-^, (iii) crRNA^-^ *cas3*^+^, (iv) crRNA^+^ *cas3*^+^. We detected robust adaptation in cells expressing *cas3* and containing the stp (crRNA^+^ *cas3*^+^). By contrast, we detected background levels of fluorescence in cells lacking *cas3* and/or the stp crRNA (Figure 1B; 0-2 YFP^+^ cells from a total of ∼80,000 each; Supplementary Dataset). Thus, we detected robust primed adaptation, but no naïve adaptation in a genetically similar strain. We cannot rule out the possibility that naïve adaptation occurs at a frequency below detection using this highly sensitive assay; nonetheless, we can conclude that primed adaptation is at least 1,000-fold more efficient than naïve adaptation in cells expressing *cas* genes at equivalent levels. Previous studies have detected naïve adaptation in *E. coli*, but these required either extremely high expression of *cas1* and *cas2* (Nuñez et al., 2014; Wang et al., 2015; Yosef et al., 2012), or bacterial growth over many days (Swarts et al., 2012). Importantly, the frequencies of primed and naïve adaptation we observe are directly comparable because *cas1* and *cas2* expression is the same in all the strains.

**Figure 1.**
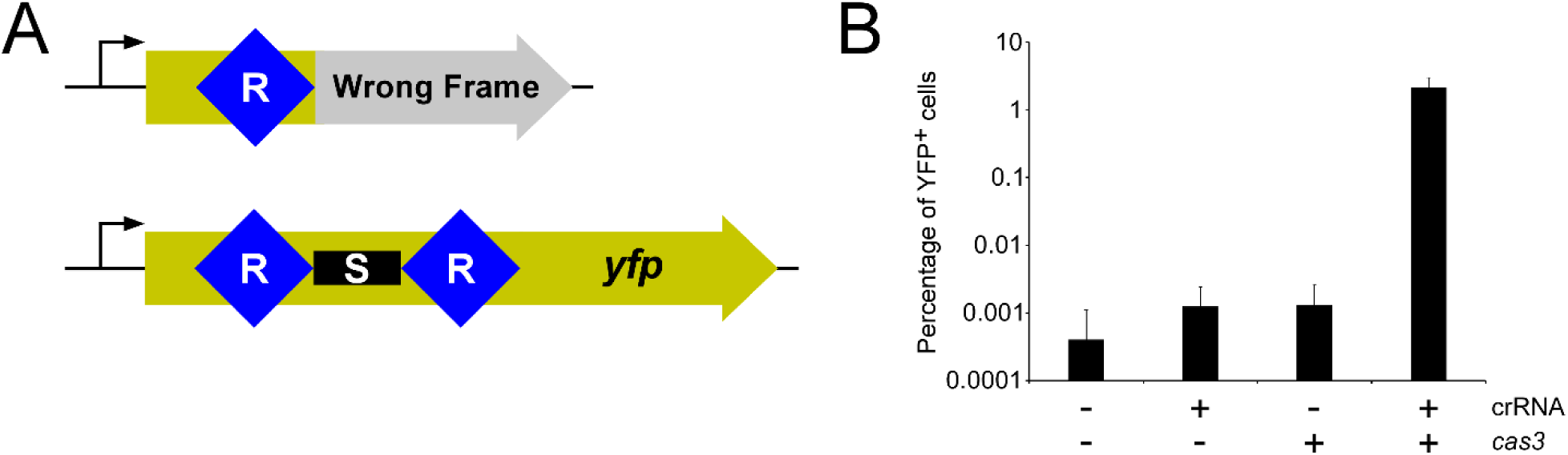
Primed adaptation is >1000-fold more efficient than naïve adaptation. **(A)** Schematic showing the *yfp* reporter construct of strain AMD688 used to quantify adaptation. The genome of the parent strain has a single CRISPR repeat embedded within a *yfp* open reading frame such that the repeat causes a frame-shift, preventing translation of YFP. Expansion of the CRISPR array by a single repeat and spacer restores the frame, permitting translation of YFP, leading to detectable fluorescence. **(B)** Percentage of YFP^+^ cells for an *E. coli* strain containing the *yfp* reporter (AMD688), either the stp (pAMD189; “crRNA +”) or an equivalent empty vector (pBAD24; “crRNA -”), and either a *cas3*-expressing plasmid (pAMD191; “cas3 +”) or an equivalent empty vector (pBAD33; “cas3 -”), following addition of arabinose to the cultures. Values plotted represent the mean of three independent biological replicates. Error bars represent one standard deviation from the mean.

Studies of other type I systems suggest that naïve adaptation is similarly inefficient, with primed adaptation being >500-fold more efficient than naïve adaptation in a type I-F system (Staals et al., 2016), and naïve adaptation being undetectable in a type I-B system (Li et al., 2014). This suggests that CRISPR immunity is extremely inefficient for prokaryotes with type I systems when encountering an invading DNA molecule that is not already at least a partial match to an existing spacer. Hence, CRISPR immunity in such a situation would likely only be effective in the context of a large, clonal population of cells.

### Mapping newly acquired spacers suggests primed adaptation can occur from chromosomal sites

Previous studies have shown that the majority of pre-spacers used during primed adaptation in *E. coli* are located on the non-target strand, and are associated with an AAG PAM (Datsenko et al., 2012; Fineran et al., 2014; Savitskaya et al., 2013; Swarts et al., 2012). To determine whether the same features are true for our experimental set-up, we induced primed adaptation using the stp, and determined the location of pre-spacers by PCR-amplification and sequencing of the expanded CRISPR arrays. We specifically looked at the first spacer sequence in CRISPR arrays that were expanded by one spacer. As expected, the majority of acquired spacers mapped to the stp (Figure 2A), and the distribution of pre-spacers on the stp was independent of the CRISPR array in which the spacer was acquired (*E*. coli has two CRISPR arrays; Figure 2B). 75.3% of pre-spacers were located on the same DNA strand as the protospacer, and 94.4% of spacers were associated with an AAG PAM. Thus, our data are consistent with earlier studies of primed adaptation.

**Figure 2.**
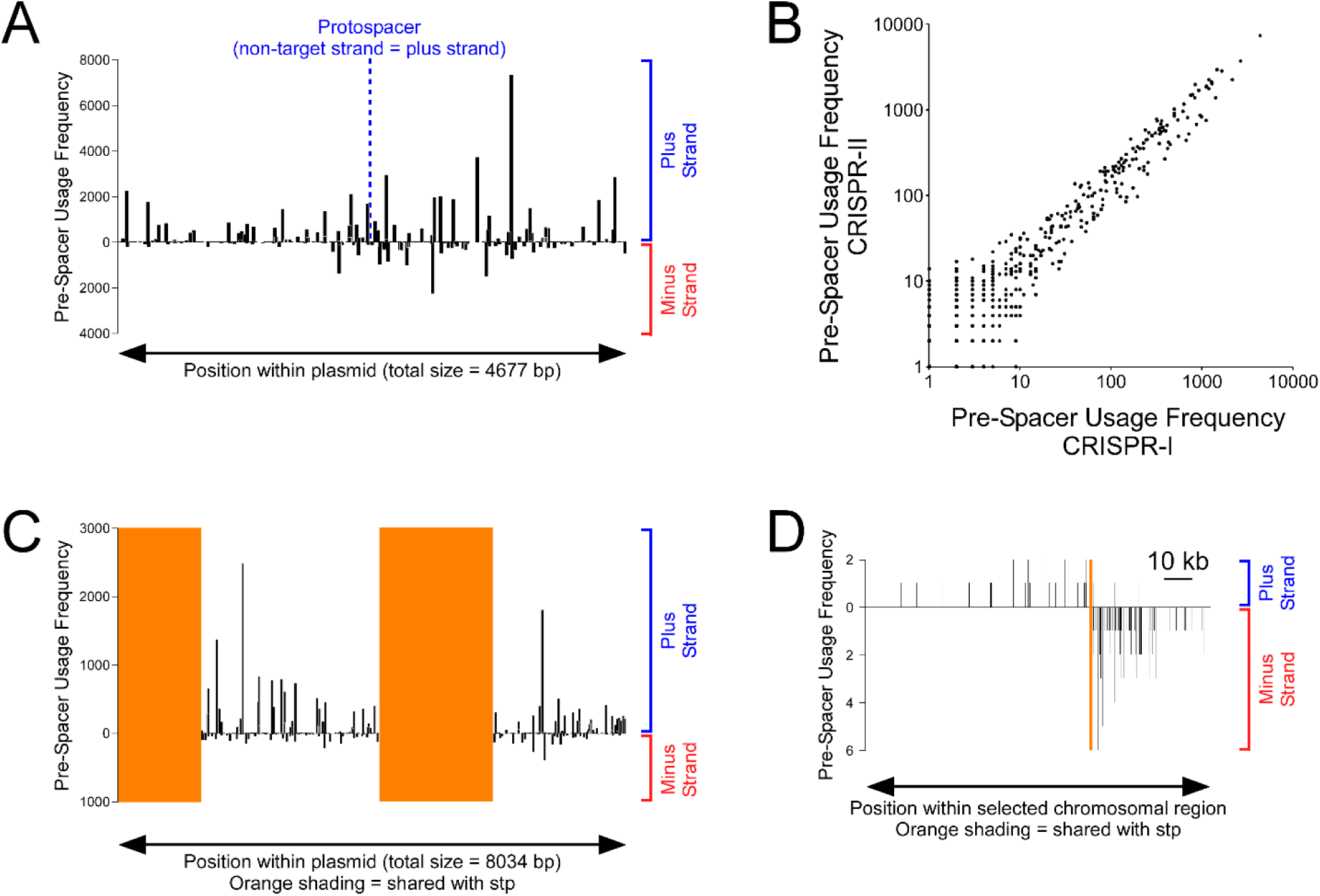
Primed adaptation from a plasmid results in primed adaptation from shared chromosomal locations. **(A)** Frequency of pre-spacer usage across the stp (pAMD189) during primed adaptation in AMD536 cells containing pAMD189 and pAMD191. Values represent the number of sequenced instances of each pre-spacer, plotted as a function of position on the plasmid. Positive values indicate pre-spacers on the forward strand; negative values indicate pre-spacers on the reverse strand. **(B)** Frequency of pre-spacer usage on the stp compared for spacers acquired in the CRISPR-I and CRISPR-II arrays. Data are only plotted for 33 bp pre-spacers that were sequenced at least once in both datasets. **(C)** Frequency of pre-spacer usage across pAMD191 during primed adaptation in AMD536 cells containing pAMD189 and pAMD191. Only pre-spacers unique to pAMD191 are shown; orange shaded regions indicate shared sequence with pAMD189. **(D)** Frequency of pre-spacer usage across a chromosomal region encompassing the *araC* gene during primed adaptation in AMD536 cells containing pAMD189 and pAMD191. Values are plotted as a function of genome position. Only pre-spacers unique to the chromosome are shown; the orange shaded region indicates shared sequence with pAMD189.

Intriguingly, many pre-spacers did not map to the stp, suggesting another source for the acquired spacers. Most of the pre-spacers that did not map to the stp mapped uniquely to the *cas3*-expressing plasmid (Figure 2C), although it is important to note that much of the sequence of this plasmid is identical to that of the stp, and reads that match shared regions were mapped to the stp. Pre-spacer usage on the *cas3*-expressing plasmid was strongly biased to one strand and to pre-spacers with an AAG PAM. Hence, we speculated that pre-spacers mapping uniquely to the *cas3*-expressing plasmid represent a second round of primed adaptation. Even though we sequenced the first acquired spacer in each array, this spacer may have been acquired as a result of primed adaptation using a newly acquired spacer in the other array. Consistent with multiple rounds of adaptation occurring in individual cells, we previously showed that primed adaptation under these conditions often leads to successive rounds of expansion of the CRISPR-II array (Cooper et al., 2018).

We also detected a small number of pre-spacers that mapped uniquely to the chromosome. We reasoned that these sequences could have been acquired due to multiple rounds of primed adaptation if the first acquired spacer matched a sequence shared between the stp and the chromosome. Consistent with this hypothesis, pre-spacers mapping uniquely to the chromosome were found adjacent to sequences shared with the stp, i.e. the *araC* gene (Figure 2D), or the transcription terminators of rRNA loci. Thus, our data indicate that primed adaptation from chromosomal pre-spacers can be readily detected using this approach.

### Primed adaptation occurs over a distance of >100 kb from the protospacer

Previous studies suggest that primed adaptation in *E. coli* occurs over relatively long distances, exceeding the size of a typical plasmid (Strotskaya et al., 2017). Given that we observed primed adaptation from chromosomal sites when targeting the stp with a crRNA, we reasoned that targeting a chromosomal site with a crRNA would lead to extensive primed adaptation from the chromosome, allowing us to infer the distance over which primed adaptation occurs. In two independent experiments, we initiated primed adaptation from protospacers located (i) immediately upstream of the *lacZ* gene, with the target strand of the protospacer on the plus strand of the genome (“*lacZ*+” protospacer), and (ii) ∼10 kb from the *lacZ* gene, inside the *mhpT* gene, with the target strand of the protospacer on the minus strand of the genome (“*mhpT*-” protospacer). Both protospacers have an AGG PAM, which can induce both interference and primed adaptation (Cooper et al., 2018; Fineran et al., 2014; Musharova et al., 2019; Xue et al., 2015). Moreover, we previously showed that targeting the *lacZ+* protospacer with a crRNA leads to robust association of Cascade (Cooper et al., 2018). Following induction of primed adaptation for each of the two protospacers, we inferred the location of pre-spacers by PCR-amplification and sequencing of the first acquired spacer in the expanded CRISPR-II array (Figure 3A; singly expanded arrays only). Thus, we observed robust primed adaptation on the chromosome, centered at each of the protospacers (Figure 3B). We presume that initiating interference and primed adaptation at chromosomal sites is lethal to *E. coli*. Consistent with this, a recent study showed that primed adaptation from a chromosomal protospacer leads to selection for deletions around the protospacer (Shiriaeva et al., 2019). Nonetheless, we were able to sequence large numbers of spacers acquired as a result of chromosomal priming, suggesting that we are capturing adaptation events as cells die.

**Figure 3.**
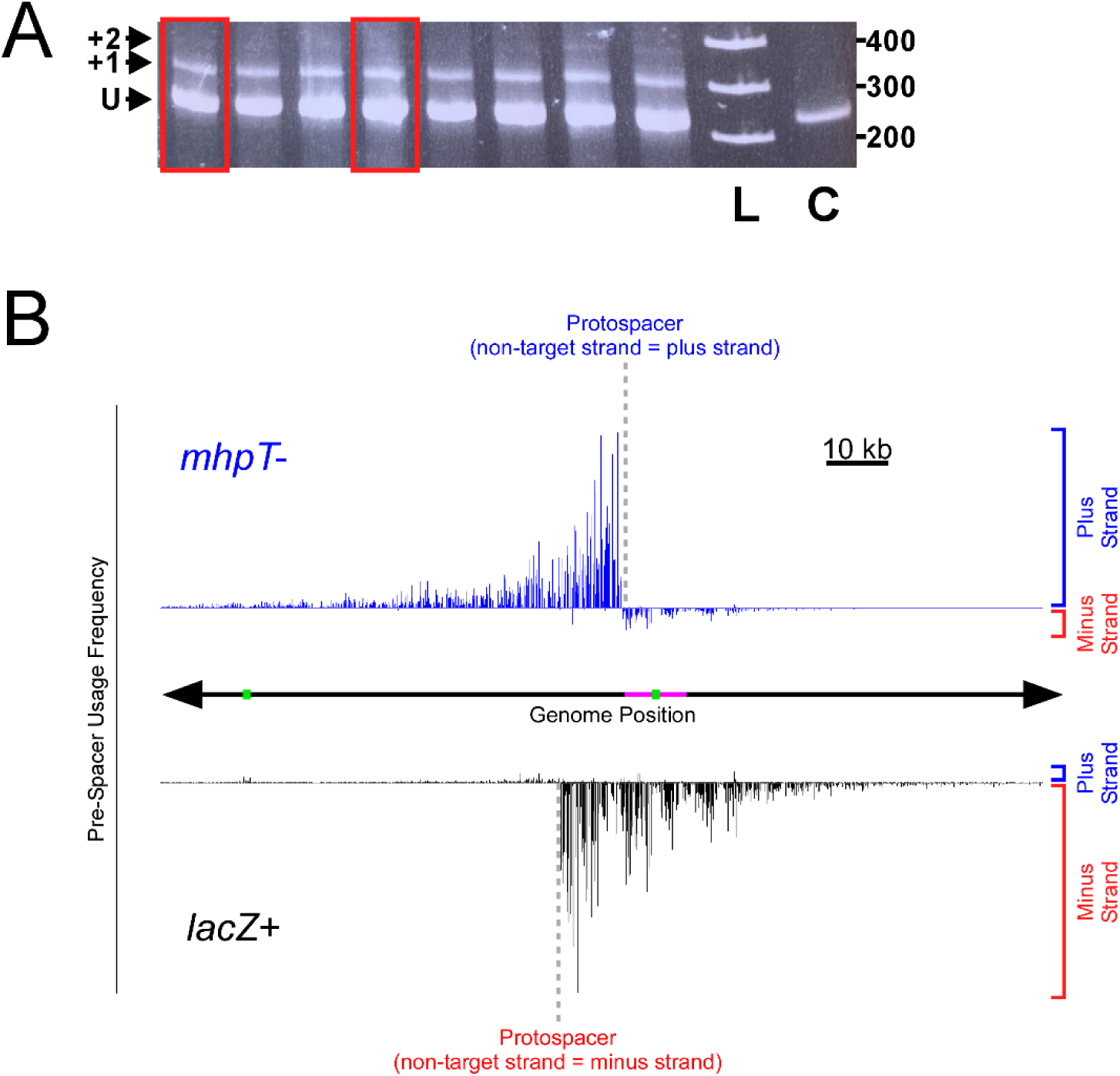
Primed adaptation on the chromosome occurs over regions of >100 kb. **(A)** Ethidium bromide-stained polyacrylamide gel showing PCR products spanning the start of the CRISPR-II array in cells undergoing primed adaptation using the *lacZ*+ protospacer. Red boxes indicate the replicate samples used for the analyses described in this work; other lanes are for samples not used in this study. The expected positions of PCR products corresponding to unexpanded (“U”), singly expanded (“+1”), and doubly expanded (“+2”) arrays are indicated on the left. “L” indicates a molecular weight ladder, with sizes of molecular weight standards (bp) indicated to the right of the gel. “C” indicates a control sample from cells not undergoing primed adaptation. **(B)** Frequency of pre-spacer usage across a chromosomal region encompassing the *lacZ*+ and *mhpT*-protospacers for AMD536 cells containing pAMD191 (encodes Cas3), and either pCB380 (encodes crRNA targeting *lacZ*+; data shown in black) or pAMD212 (encodes crRNA targeting *mhpT*-; data shown in blue). Values plotted on the *y*-axis represent the relative number of sequenced instances of each pre-spacer, plotted as a function of genome position (*x*-axis). Positive values indicate pre-spacers on the forward strand; negative values indicate pre-spacers on the reverse strand. Green boxes indicate positions of repetitive insertion element sequences. The pink box indicates the region analyzed in Figure 6B.

Consistent with primed adaptation studies using plasmids, the majority of chromosomal pre-spacers were located on the non-target strand, upstream of the protospacer (Figure 3B). Remarkably, we detected primed adaptation over a distance of >100 kb upstream of each of the two protospacers, with the extent of primed adaptation decreasing as a function of distance from the protospacer, albeit with considerable local variation (Figure 3B). Of note, recent studies described priming on the *E. coli* chromosome over a similar distance (Shiriaeva et al., 2019), and unidirectional deletions over distances up to 100 kb upon expression of type I-E CRISPR-Cas systems in human cells (Dolan et al., 2019; Morisaka et al., 2019). Our data strongly suggest that Cas3 can translocate >100 kb from the protospacer during primed adaptation. Since the strain we used has both CRISPR arrays intact, it is possible that some spacers acquired in CRISPR-II represent two rounds of primed adaptation, with the first round of primed adaptation leading to spacer acquisition in CRISPR-I. However, we believe this is a rare occurrence, for two reasons. First, most cells had an unexpanded CRISPR-II array, and we observed very few doubly expanded arrays (Figure 3A), although the PCR assay is not sensitive enough to detect low levels of doubly expanded arrays. Second, there is an insertion element ∼15 kb away from the *lacZ*+ protospacer that contains many frequently used pre-spacers. This insertion element sequence is duplicated at several other genomic locations, but we observed very few pre-spacers adjacent to the duplicated regions. Hence, we conclude that very few spacers acquired in CRISPR-I from this insertion element led to a second round of primed adaptation in CRISPR-II. Nonetheless, even if a substantial proportion of the pre-spacers we observed result from two rounds of primed adaptation, a single round of primed adaptation must still be able to occur over a distance of >50 kb.

The large distance over which we observed primed adaptation explains why the efficiency of primed adaptation from a plasmid for type I-E CRISPR-Cas systems does not appear to decrease as a function of distance from the protospacer (Savitskaya et al., 2013); presumably, Cas3 translocates many times around the circular plasmid. It also suggests that primed adaptation would facilitate acquisition of spacers from any region of even large bacteriophage genomes or plasmids, which may increase the effectiveness of the immune response. The large distance over which primed adaptation occurs on the chromosome also means that there are many more unique pre-spacers than used during primed adaptation from a plasmid. We reasoned that this much larger set of pre-spacers would allow for a more in-depth analysis of the pre-spacer features that influence the efficiency of primed adaptation.

### Off-target Cascade binding sites are not associated with primed adaptation

We previously showed that Cascade association with a DNA target requires only limited base-pairing between the crRNA spacer and the protospacer. Hence, Cascade binds to many “off-target” chromosomal sites that typically have between 5 and 10 nt matches to the 5’ end of a crRNA spacer, immediately flanked by an optimal (AAG) PAM (Cooper et al., 2018). When targeting the *lacZ*+ protospacer with a crRNA, we observed Cascade binding to ∼100 off-target sites (Cooper et al., 2018). We determined the level of primed adaptation from the non-target strand in the 10 kb upstream each of the 76 off-target sites associated with a seed sequence resembling that of the on-target *lacZ*+ protospacer. The level of Cascade association and the local pre-spacer usage frequency were not significantly correlated (Figure 4; Spearman’s Correlation Test, two-tailed, *p* = 0.72), with pre-spacer usage near off-target sites typically being due to duplicated sequences with identical copies close to the *lacZ*+ protospacer. Moreover, the local pre-spacer usage frequency in regions adjacent to off-target Cascade binding sites was not significantly higher than that of 1000 randomly selected genomic locations (Mann-Whitney U Test *p* = 0.054). Thus, our data suggest that off-target Cascade binding events do not lead to primed adaptation, consistent with our previous observation that extensive PAM-distal mismatches between a crRNA spacer and its cognate protospacer prevent primed adaptation (Cooper et al., 2018).

**Figure 4.**
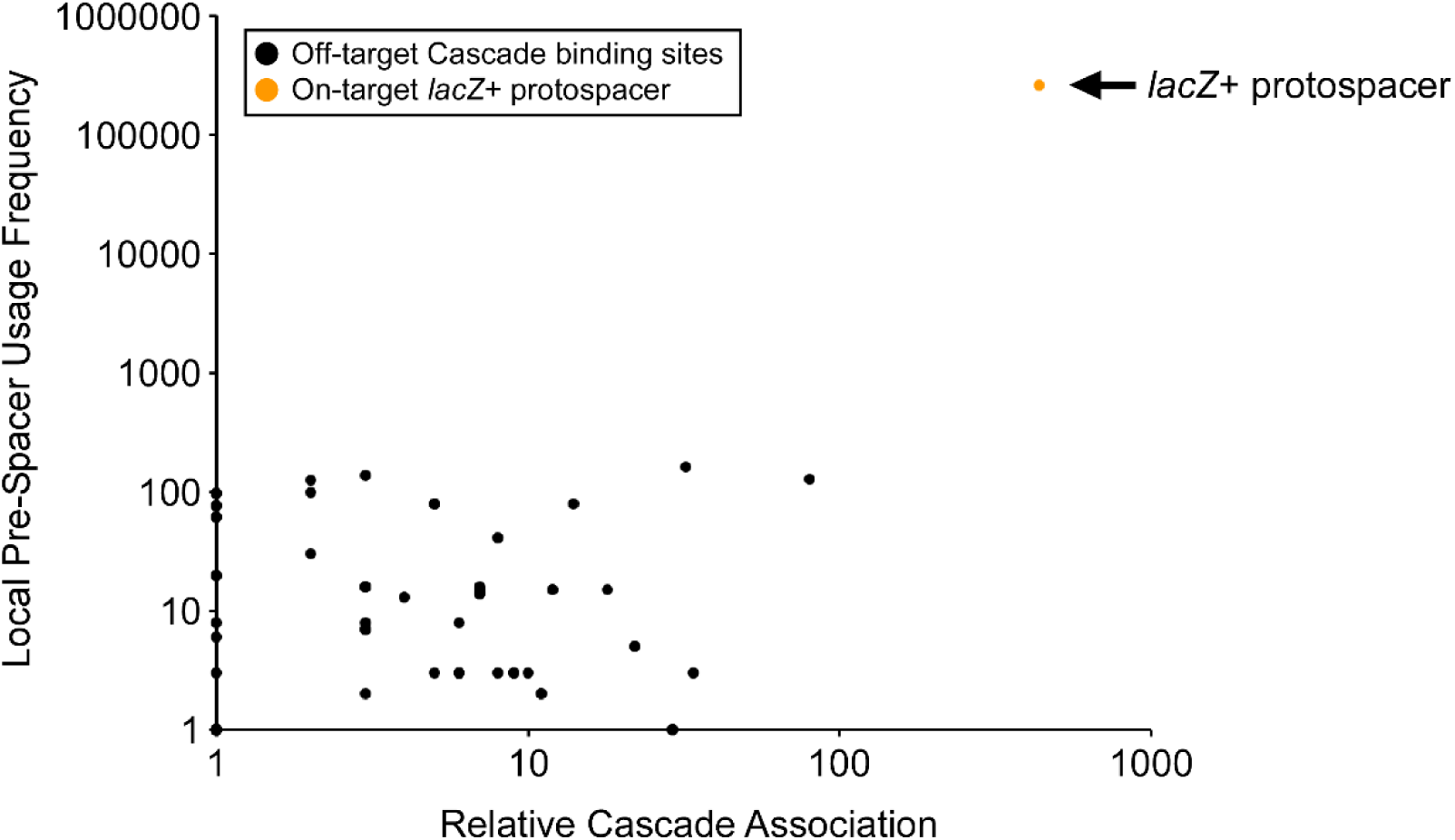
Off-target Cascade binding sites are not associated with detectable primed adaptation. Frequency of pre-spacer usage in the 10 kb upstream of the on-target *lacZ*+ protospacer (orange datapoint), or off-target protospacers identified by (Cooper et al., 2018), on the non-target strand, only counting 33 bp pre-spacers. Relative Cascade association with each target sequence, as determined by ChIP-seq (Cooper et al., 2018), is shown on the *x*-axis.

### Uneven distribution of chromosomal pre-spacers relative to the protospacer

The vast majority of pre-spacers we identified are 33 bp long (98.6% for both the *lacZ*+ and *mhpT*-protospacers; note that pre-spacers include the last base of the PAM, which is known to be added to the CRISPR array by Cas1-Cas2; (Goren et al., 2012)). Unless specifically stated, the analyses described below focus exclusively on the 33 bp pre-spacers. We also excluded genomic regions covered by insertion elements, repetitive sequences that confound alignment of DNA sequence reads. Lastly, since the independent replicate datasets for each protospacer were highly similar (Figure 5A + B), all analyses described below use datasets where the two replicates were combined.

**Figure 5.**
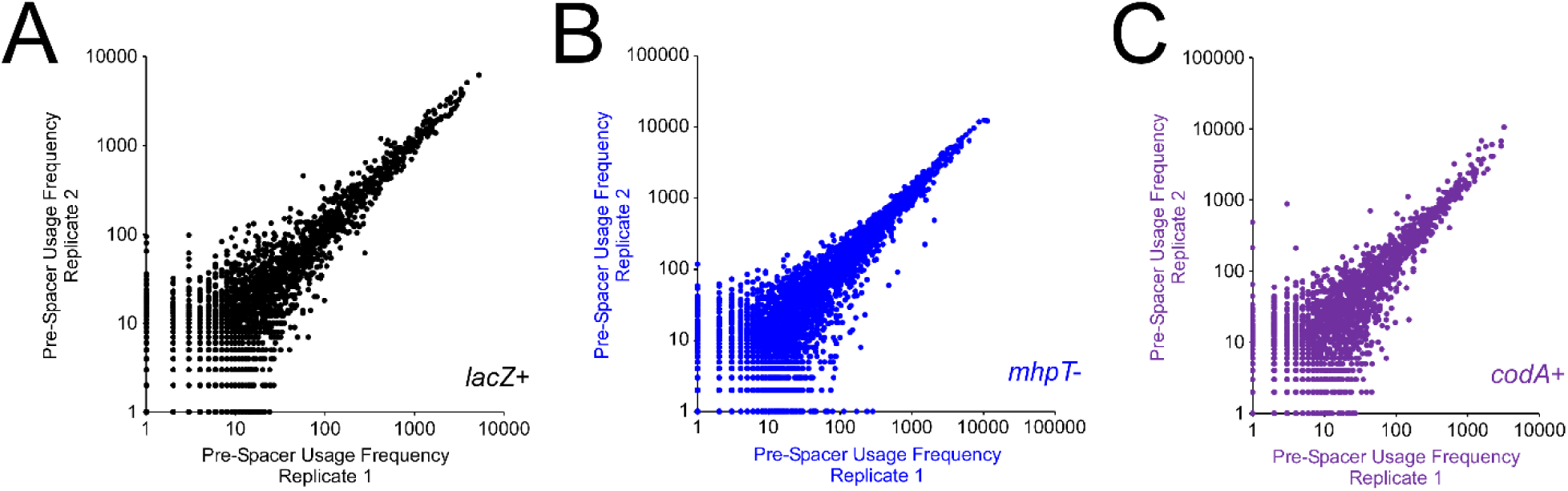
Replicate primed adaptation datasets for chromosomal protospacers are highly reproducible. Frequency of pre-spacer usage across the chromosome for each of two replicates for AMD536 cells containing pAMD191 (encodes Cas3), and pCB380 (encodes crRNA targeting *lacZ*+), **(B)** pAMD212 (encodes crRNA targeting *mhpT*-), or **(C)** pAMD211 (encodes crRNA targeting *codA*+). Values represent the number of sequenced instances of each pre-spacer, for every 33 bp pre-spacer that mapped to the chromosome.

Although the majority of pre-spacers were located on the non-target strand, upstream of the protospacers, we also observed a low level of primed adaptation on the target strand, upstream of the protospacer (Figure 3 + 6A). As for pre-spacers on the non-target strand, the frequency of pre-spacer usage on the target-strand decreased as a function of distance from the protospacer, albeit with considerable local variation. Strikingly, the fraction of pre-spacers on the target strand, upstream of the protospacer, was ∼4-fold higher for the *mhpT*-protospacer than for the *lacZ*+ protospacer, suggesting that the protospacer sequence context determines the bias in primed adaptation towards the non-target strand (Figure 3 + 6A). Since the two protospacers we used are only ∼10 kb apart, and are in opposite DNA orientations, many of the *target* strand pre-spacers upstream of the *mhpT*-protospacer are from the same genomic region and strand as *non-target* strand pre-spacers upstream of the *lacZ*+ protospacer (pink box in Figure 3). Indeed, we observed very similar profiles of pre-spacer location and abundance across this region for the two datasets (Figure 6B), indicating that primed adaptation on the target strand upstream of the protospacer likely occurs by the same mechanism as primed adaptation on the non-target strand upstream of the protospacer.

**Figure 6.**
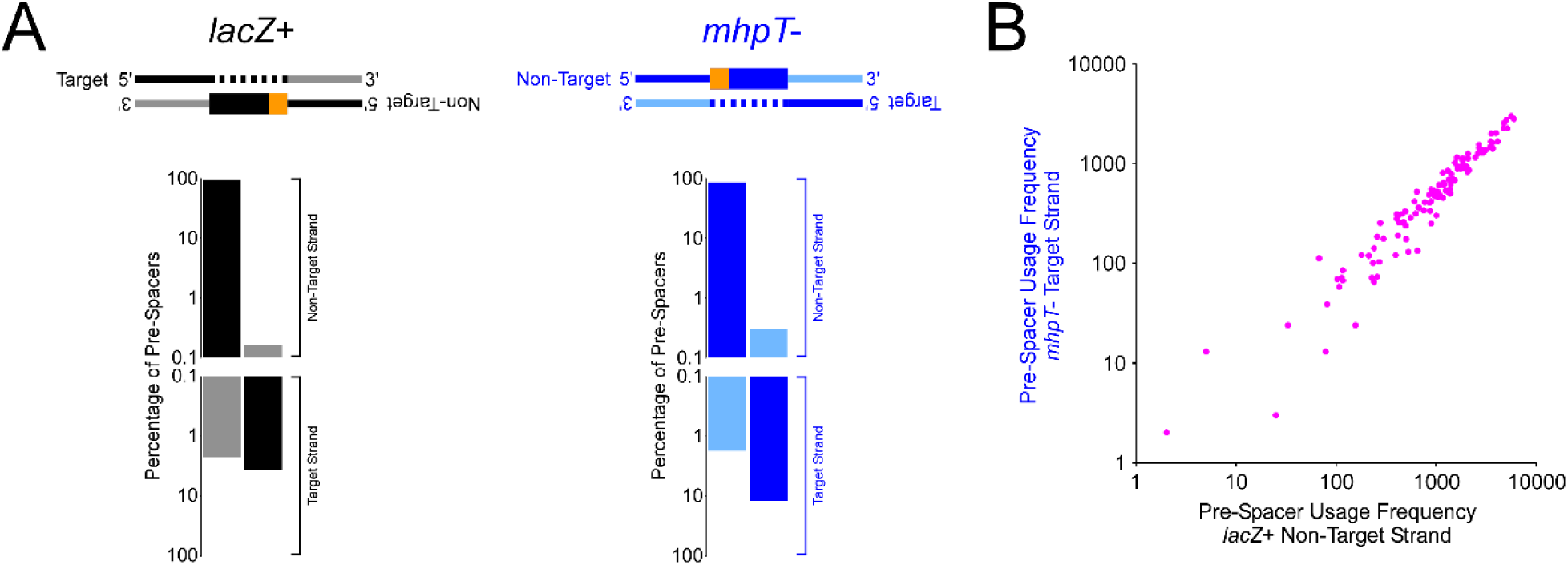
The majority of primed adaptation occurs on the non-target strand, upstream of the protospacer. **(A)** Distribution of pre-spacer usage for *lacZ*+ (black) and *mhpT*- (blue) between the non-target (positive values) and target (negative values) strands, and between the upstream (dark colors) and downstream (light colors) directions. **(B)** Frequency of pre-spacer usage in the 10 kb upstream of the *mhpT*-protospacer on the target strand for AMD536 cells containing pAMD191 (encodes Cas3), and pCB380 (encodes crRNA targeting *lacZ*+; *x*-axis) or pAMD212 (encodes crRNA targeting *mhpT*-; *y*-axis). Values represent the number of sequenced instances of each pre-spacer, for every 33 bp pre-spacer associated with an AAG PAM.

The strong bias in pre-spacer usage to the region on the non-target strand, upstream of the protospacer, presumably reflects a bias in the direction of Cas3 translocation. We propose that Cas3 translocation is strongly biased to the non-target strand because of the inherent asymmetry in the Cascade-bound protospacer. In type I-F systems, the only characterized type I systems where Cas3 translocation appears to occur with equal efficiency on both the target and non-target strands, Cas2 is fused to Cas3, such that there are two Cas3 subunits in the Cas1-Cas2-Cas3 complex (Fagerlund et al., 2017; Rollins et al., 2017). We suggest that this allows translocation of Cas2-3 in either strand with roughly equal efficiency, presumably using a different Cas3 subunit for each of the two strands. However, there are likely to be other factors that influence the degree to which Cas3 selects strand, since our data show that the ratio of target:non-target strand usage varies between protospacers (Figure 6A). Consistent with this idea, a recent study of primed adaptation from a chromosomal protospacer in *E. coli* showed a much higher level of pre-spacer usage on the target strand, upstream of the protospacer (Shiriaeva et al., 2019).

### Acquisition of spacers with non-AAG PAMs is frequently due to slipping

We examined all pre-spacers associated with the *lacZ*+ and *mhpT*-protospacers on the non-target strand, in the 10 kb region upstream of the protospacer. 95.0% and 96.0% of pre-spacer usage was associated with an AAG PAM for the *lacZ*+ and *mhpT*-protospacers, respectively. While the large majority of pre-spacer usage is associated with an AAG PAM, we were interested to determine the basis for selecting pre-spacers with a non-AAG PAM. Previous studies of type I-E, type I-B, type I-C and type I-F CRISPR-Cas systems have shown that pre-spacers with non-canonical PAMs can be acquired due to a phenomenon known as “slipping”, whereby the pre-spacer with a non-optimal PAM is positioned one or two nucleotides from a canonical, optimal PAM-associated pre-spacer (Li et al., 2017; Rao et al., 2017; Shmakov et al., 2014; Staals et al., 2016). To determine the frequency of slipping in our data, we calculated all pairwise distances for non-AAG PAM pre-spacers to AAG PAM pre-spacers on the non-target strand, in the 10 kb upstream of each of the protospacers. We observed a strong enrichment of pre-spacers that could be assigned to “-1”, “+1”, or “+2” slips (Figure 7A + B), with +1 slips being the most frequent, accounting for 37% of all non-AAG PAM pre-spacer usage. Our data indicate that together, +1 slips and -1 slips represent the majority (60%) of primed spacer acquisition on the non-target strand for pre-spacers not associated with an AAG PAM. The mechanism of slipping is not known, but may be due to imprecise processing of Cas1-Cas2-bound pre-spacer DNA by exonucleases (Kim et al., 2020; Ramachandran et al., 2020). The remaining pre-spacers that lack an AAG PAM and are not associated with slipping events typically have suboptimal PAMs, with the third base of the PAM strongly enriched for G, and the second base moderately enriched for A or C (Figure 7C). Thus, our data suggest that while the specificity for an AAG PAM is high during primed adaptation, there is a hierarchy of selectivity within the PAM, with the third position being the most important, and the first position being the least important. It is possible that this hierarchy influences the relative frequency of the different types of slipping events.

**Figure 7.**
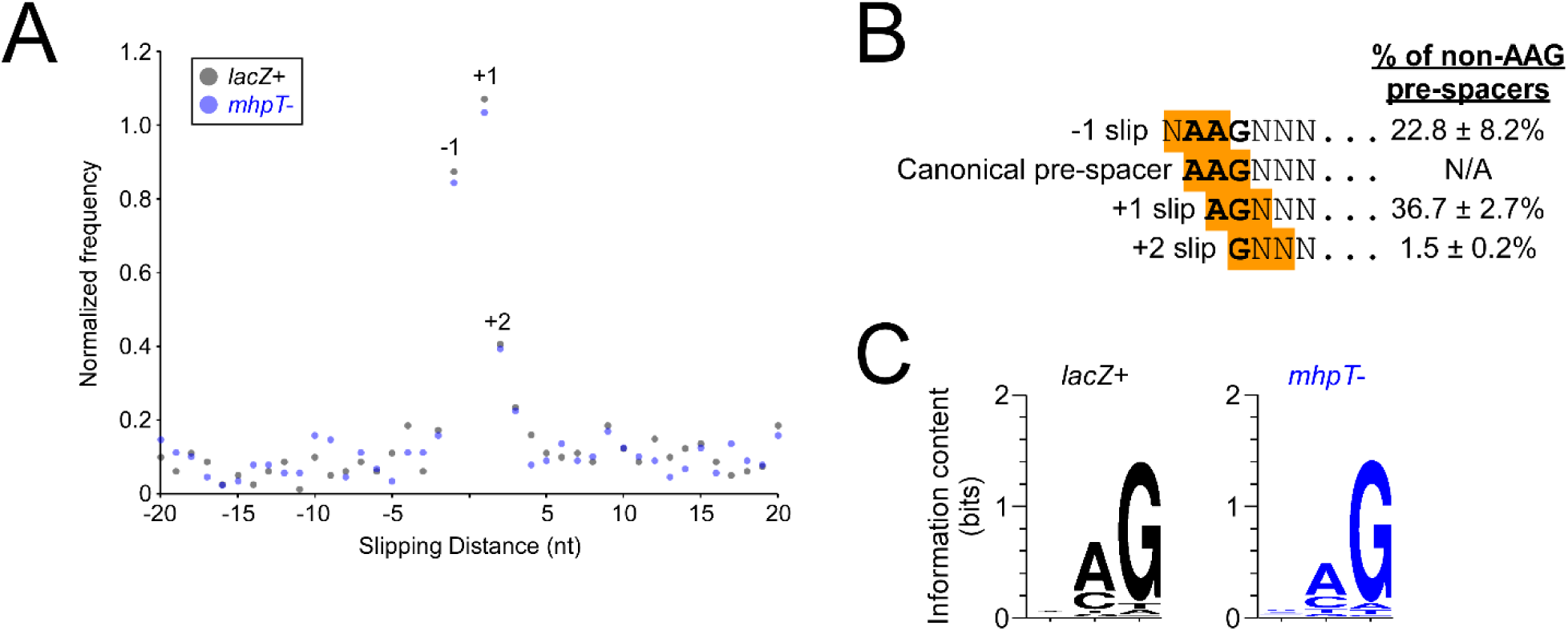
Slipping accounts for the majority of pre-spacers on the non-target strand that are not associated with an AAG PAM. **(A)** Normalized frequency of distances (“slipping distance”; *x*-axis) between uniquely positioned protospacers with an AAG PAM and uniquely positioned protospacers with a non-AAG PAM, for the 10 kb upstream of the *lacZ*+ (black datapoints) and *mhpT*- (blue datapoints) protospacers, on the non-target strand. Values were normalized to the total number of pairwise comparisons. **(B)** Distribution of pre-spacer usage for all pre-spacers with a non-AAG PAM, for the 10 kb upstream of the protospacer on the non-target strand. Values represent averages from the *lacZ*+ and *mhpT*-experiments, with error values representing one standard deviation from the mean. **(C)** DNA sequence logos for PAMs from pre-spacers with a non-AAG PAM, for the 10 kb upstream of each of the *lacZ*+ and *mhpT*-protospacers on the non-target strand, excluding pre-spacers associated with -2, -1, +1 or +2 slipping events. Logos were generated from all pre-spacer usage (i.e. not just unique pre-spacerse).

Analysis of slipping in a type I-F CRISPR-Cas system demonstrated that slipped pre-spacers are associated with PAMs that lead to efficient primed adaptation (Jackson et al., 2019). Our data indicate that the most common form of slipping is a +1 slip, which would lead to an AGN PAM (Figure 7B). The next most common form of slipping is a -1 slip, which would lead to a NAA PAM (Figure 7B). Previous studies of PAM sequences that lead to interference and/or primed adaptation indicate that of all the possible AGN and NAA PAMs, only AGT, and possibly CAA, fail to lead to interference and/or primed adaptation (Musharova et al., 2019; Xue et al., 2015). Thus, slipping in the *E. coli* CRISPR-Cas system likely leads to acquisition of a functional spacer in most cases.

Previous studies of naïve adaptation showed only ∼25% of pre-spacer usage in *E. coli* is associated with AAG PAMs (Levy et al., 2015; Yosef et al., 2013), in contrast to the >95% observed for primed adaptation in our study and other studies (Savitskaya et al., 2013). One possible explanation is that the high level of Cas1-Cas2 overexpression required to detect naïve adaptation causes a reduction in PAM specificity. An alternative hypothesis is that Cas1-Cas2 association with the translocating Cas3-containing complex during priming leads to an increase in specificity for AAG; Cas3 cuts DNA with some sequence specificity (Künne et al., 2016), although this low level of specificity alone is unlikely to account for the large difference in PAM specificity between naïve and primed adaptation. Cas8e, a subunit of Cascade, binds preferentially to AAG PAMs (Hayes et al., 2016; Westra et al., 2013) (note that Cas8e interacts primarily with the CTT on the opposite strand), albeit with considerable flexibility in sequence preference (Musharova et al., 2019; Xue et al., 2015). It is possible that Cas8e translocates with Cas3 and facilitates nicking at AAG sequences. In support of this idea, Cas8 is fused to Cas3 in some bacterial species (Westra et al., 2012). Moreover, Cas8e associates only weakly with the rest of the Cascade complex, and frequently dissociates from the complex *in vitro* (Jore et al., 2011), in particular when there are suboptimal bases in the PAM or in the PAM-proximal region of the protospacer (Jung et al., 2017). Cas8e has also been suggested to dissociate from the Cascade complex *in vivo*, based on single-molecule imaging data (Vink et al., 2020). However, *in vitro* reconstitution of Cas3 translocation for the *Thermobifida fusca* Cas3 suggests that translocating Cas3 does not associate with Cas8e (Dillard et al., 2018).

### Acquisition of spacers from the target strand is frequently due to flipping, often following a slipping event

We examined all pre-spacers associated with the *lacZ*+ and *mhpT*-protospacers on the target strand, in the 10 kb downstream of the protospacer, i.e. on the opposite strand to where the most frequently used pre-spacers are located. 2.4% and 2.1% of pre-spacer usage within 10 kb of the protospacer, on this side (i.e. upstream of the protospacer on the non-target strand, or downstream of the protospacer on the target strand) is on the target strand for the *lacZ*+ and *mhpT*-protospacers, respectively. While the large majority of pre-spacer usage is on the non-target strand, we were interested to determine the basis for selecting pre-spacers from the target strand. Previous studies of type I-E, type I-B, type I-C and type I-F CRISPR-Cas systems have shown that pre-spacers with non-canonical PAMs can be acquired due to a phenomenon known as “flipping”, whereby a pre-spacer is selected from the non-target strand but then inserted into the CRISPR array in the opposite orientation (Figure 8A) (Li et al., 2017; Rao et al., 2017; Shmakov et al., 2014; Staals et al., 2016). To determine the frequency of flipping in our data, we calculated all pairwise distances for target-strand pre-spacers to AAG pre-spacers on the non-target strand, in the 10 kb upstream of each of the protospacers. We reasoned that a distance of 32 bp would represent a flipping event involving a canonical pre-spacer with an AAG PAM. We observed an enrichment of distances that support flipping events involving spacers with a -1, +1, or +2 nt slip, in addition to flipping events involving non-slipped pre-spacers (Figure 8B). Thus, flipping occurs more frequently with slipped pre-spacers than with non-slipped pre-spacers. We speculate that pre-spacers from the non-target strand with an AAG PAM are unlikely to be inserted into the CRISPR array in the incorrect orientation, whereas pre-spacers generated by slipping events are more likely to be flipped because they have a suboptimal PAM.

**Figure 8.**
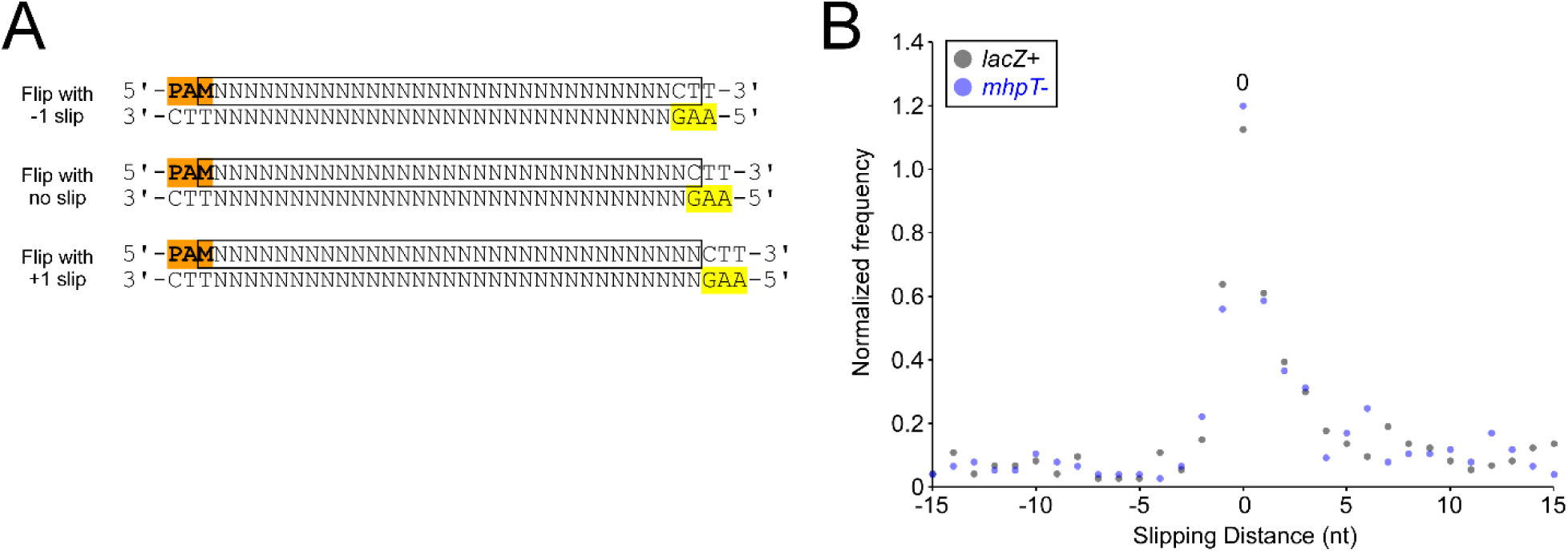
Flipping often occurs following a slipping event. **(A)** Schematic showing different possible flipping events. The yellow highlighted sequences indicate the AAG PAM associated with the non-flipped pre-spacer. **(B)** Normalized frequency of distances (“slipping distance”; *x*-axis) between uniquely positioned protospacers with an AAG PAM on the non-target strand, and uniquely positioned protospacers on the target strand, for the 10 kb upstream of the *lacZ*+ (black datapoints) and *mhpT*- (blue datapoints) protospacers on the non-target strand (i.e. 10 kb downstream of the protospacers on the target strand). Values were normalized to the total number of pairwise comparisons.

### Pre-spacers with non-canonical lengths are often associated with slipping

While the large majority of pre-spacers are 33 bp long, we were interested to determine the basis for selecting pre-spacers with non-canonical lengths. 0.3% of both *lacZ*+ and *mhpT*-pre-spacer usage was associated with pre-spacers that are 32 bp long, and 1.1% of pre-spacer usage for both *lacZ*+ and *mhpT*-pre-spacers was associated with pre-spacers that are 34 bp long. We chose to focus on the region <10 kb upstream of the protospacer, on the non-target strand, where the usage of pre-spacers of non-canonical lengths is highest (89.8% and 82.9% of all non-canonical length pre-spacers for *lacZ*+ and *mhpT*-protospacers, respectively). The majority of 32 bp pre-spacer usage for both *lacZ*+ and *mhpT*-was associated with AAG PAMs; hence, these pre-spacers are equivalent to efficiently used 33 bp pre-spacers, but lack the most PAM-distal base. These AAG-associated pre-spacers account for 87.7% (lacZ+) and 95.7% (*mhpT*-) of all 32 bp pre-spacer usage. Similarly, the majority of 34 bp pre-spacer usage was associated with AAG PAMs; hence, these pre-spacers are equivalent to efficiently used 33 bp pre-spacers, but with an additional base at the PAM-distal end. These AAG-associated pre-spacers account for 79.7% (*lacZ*+) and 86.7% (*mhpT*-) of all 34 bp pre-spacer usage. There is also a significant enrichment for 32 bp pre-spacer usage associated with +1 slips, and for 34 bp pre-spacer usage associated with -1 slips (Figure 9), indicating that slipping sometimes involves maintaining the PAM-distal boundary while either shortening or lengthening the spacer. The equivalent observation has been made for type I-B, type I-C and type I-F CRISPR-Cas systems (Li et al., 2017; Rao et al., 2017; Staals et al., 2016).

**Figure 9.**
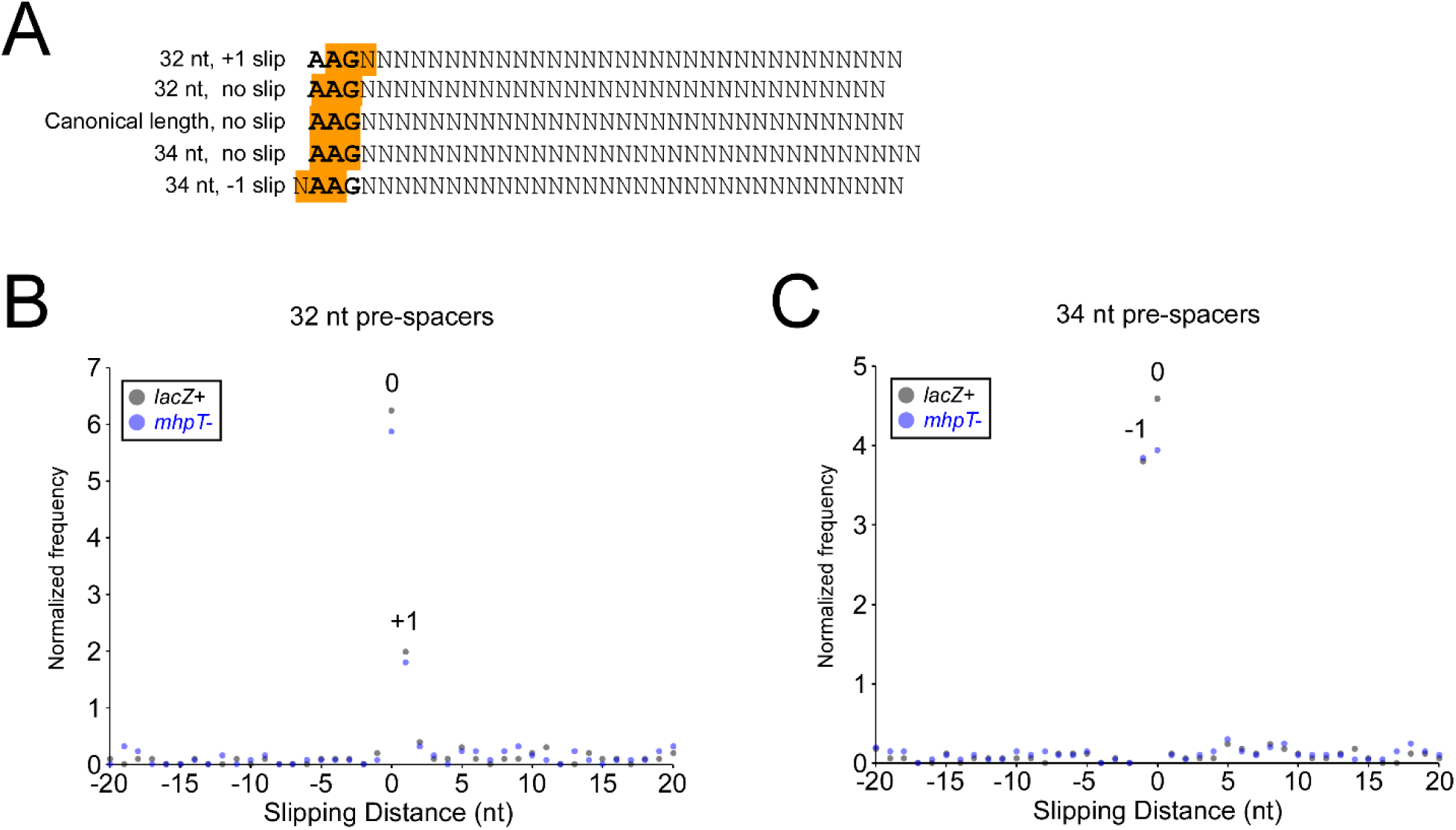
Pre-spacers with non-canonical lengths are often associated with slipping. **(A)** Schematic showing pre-spacers of non-canonical lengths associated with an AAG PAM in either a slipped or non-slipped configuration. **(B)** Normalized frequency of distances (“slipping distance”; *x*-axis) between uniquely positioned, 33 bp protospacers with an AAG PAM and uniquely positioned 32 bp protospacers, or **(C)** 33 bp protospacers, for the 10 kb upstream of the *lacZ*+ (black datapoints) and *mhpT*- (blue datapoints) protospacers, on the non-target strand. Values were normalized to the total number of pairwise comparisons.

### The efficiency of primed adaptation is reduced for pre-spacers within 200 bp of the protospacer

To determine if the pattern of pre-spacer usage is dependent upon the identity of the protospacer, we initiated primed adaptation by targeting a location ∼10 kb downstream of the *lacZ*+ protospacer, within the *codA* gene; we refer to this as the *codA*+ protospacer. Replicate datasets for *codA*+ were highly correlated (Figure 5C); hence we combined the two datasets for further analysis. As expected, we detected pre-spacer usage over a ∼100 kb distance from the *codA*+ protospacer, with the majority of pre-spacer positions overlapping those used when targeting the *lacZ*+ protospacer (Figure 10A). We compared the frequency of pre-spacer usage for pre-spacers <10 kb upstream of the *lacZ*+ protospacer on the non-target strand when targeting either the *lacZ*+ or *codA*+ protospacer. In almost all cases, the relative usage of pre-spacers was similar for each of the protospacers targeted (Figure 10B), strongly suggesting that pre-spacer usage frequency is an inherent property of the pre-spacer, and is not related to the identity of the protospacer. However, the three pre-spacers located within 200 bp of the *lacZ*+ protospacer were used proportionally less frequently when targeting the *lacZ*+ protospacer than when targeting the *codA*+ protospacer (Figure 10B). Thus, our data suggest that pre-spacers very close to the protospacer are used inefficiently. Biochemical studies of Cas3 suggested that Cas3 generates a single-stranded region adjacent to the Cascade-bound protospacer of <300 nt (Redding et al., 2015) or <500 nt (Dillard et al., 2018), for Cas3 from *E. coli* or *T. fusca*, respectively. We propose that Cas3 generates a single-stranded region of a similar length *in vivo*, and that no spacers can be acquired from the single-stranded DNA. Our data suggest that in the region <200 bp from the protospacer, the relative efficiency of pre-spacer usage frequency increases as a function of increasing distance from the protospacer (Figure 10B). Hence, we propose that Cas3 transitions stochastically between a state where it nicks DNA frequently, generating an extended region of single-stranded DNA, and a state where it nicks DNA infrequently, with the average distance traveled until the transition being 100-200 bp.

**Figure 10.**
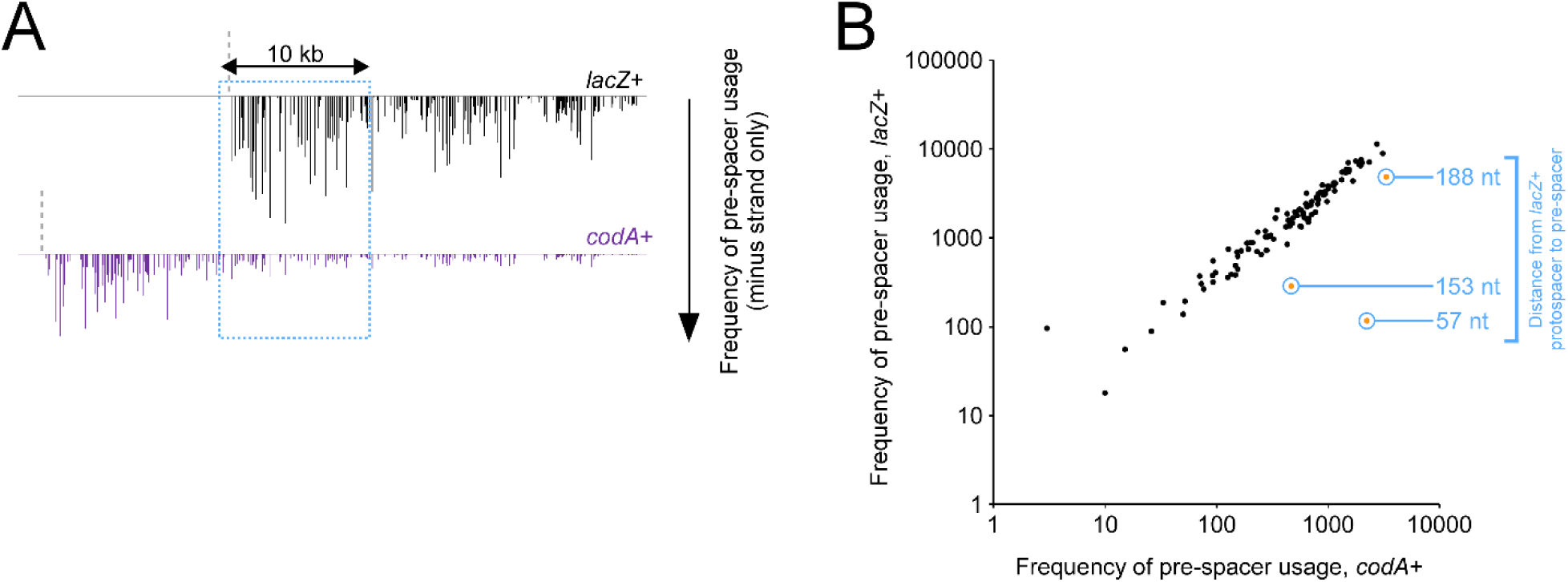
Pre-spacers with 200 nt of the protospacer are used less frequently. **(A)** Frequency of pre-spacer usage across a chromosomal region encompassing the *lacZ*+ and *codA*+ protospacers for AMD536 cells containing pAMD191 (encodes Cas3), and either pCB380 (encodes crRNA targeting *lacZ*+; data shown in black) or pAMD211 (encodes crRNA targeting *codA*+; data shown in purple). Values represent the relative number of sequenced instances of each pre-spacer, plotted as a function of genome position. Data are only plotted for pre-spacers on the reverse strand. The blue box indicates the 10 kb region analyzed in panel B. The vertical dashed lines indicate the positions of the protospacers. **(B)** Frequency of pre-spacer usage in the 10 kb upstream of the *lacZ*+ protospacer on the non-target strand for AMD536 cells containing pAMD191 (encodes Cas3), and pAMD211 (encodes crRNA targeting *codA*+; *x*-axis) or pCB380 (encodes crRNA targeting *lacZ*+; *y*-axis). Values represent the number of sequenced instances of each pre-spacer, for every 33 bp pre-spacer associated with an AAG PAM. Data for pre-spacers within 200 nt of the *lacZ*+ protospacer are circled, and the distance from the *lacZ*+ protospacer is indicated.

### The efficiency of primed adaptation is reduced for pre-spacers with an internal AAG

It is well established that the frequency of pre-spacer usage varies considerably between nearby pre-spacer sequences (Musharova et al., 2017, 2018; Savitskaya et al., 2013). However, the basis for this variability is poorly understood. As described above, the frequency of pre-spacer usage decreases globally as a function of increasing distance from the protospacer (Figure 3 + 11A+B). We fitted these data to an exponential decay model, relating pre-spacer usage to distance from the protospacer. We calculated the ratio of the observed usage frequency to the modeled usage frequency for each pre-spacer. Importantly, there was no significant difference in the observed:expected usage frequency ratios for pre-spacers within the essential genes *secD, secF, ribD, ribE* and *thiL* and those for all other pre-spacers within 100 kb of the *lacZ*+ protospacer (One-tailed Mann Whitney U test *p* = 0.55), indicating that there is no selection against pre-spacers in essential genes. We then selected “enriched” pre-spacers that have a >4-fold higher usage frequency than predicted by the model, and “de-enriched” pre-spacers that have a >4-fold lower usage frequency than predicted by the model. We determined the frequency of all tetranucleotide and trinucleotide sequences within the sets of enriched and de-enriched pre-spacers (Figure 11C-D). Almost all tetranucleotide and trinucleotide sequences were found at similar frequencies within the enriched and de-enriched pre-spacers. However, AAG-containing pre-spacers were found far less frequently in enriched pre-spacers than in de-enriched pre-spacers. We compared the frequency of pre-spacer usage for AAG-containing and non-AAG-containing pre-spacers (Figure 11A-B). The frequency was, on average, ∼3.5-fold higher for non-AAG-containing pre-spacers than for AAG-containing pre-spacers. We conclude that the presence of an AAG within a pre-spacer substantially reduces the frequency with which that pre-spacer is selected during primed adaptation.

**Figure 11.**
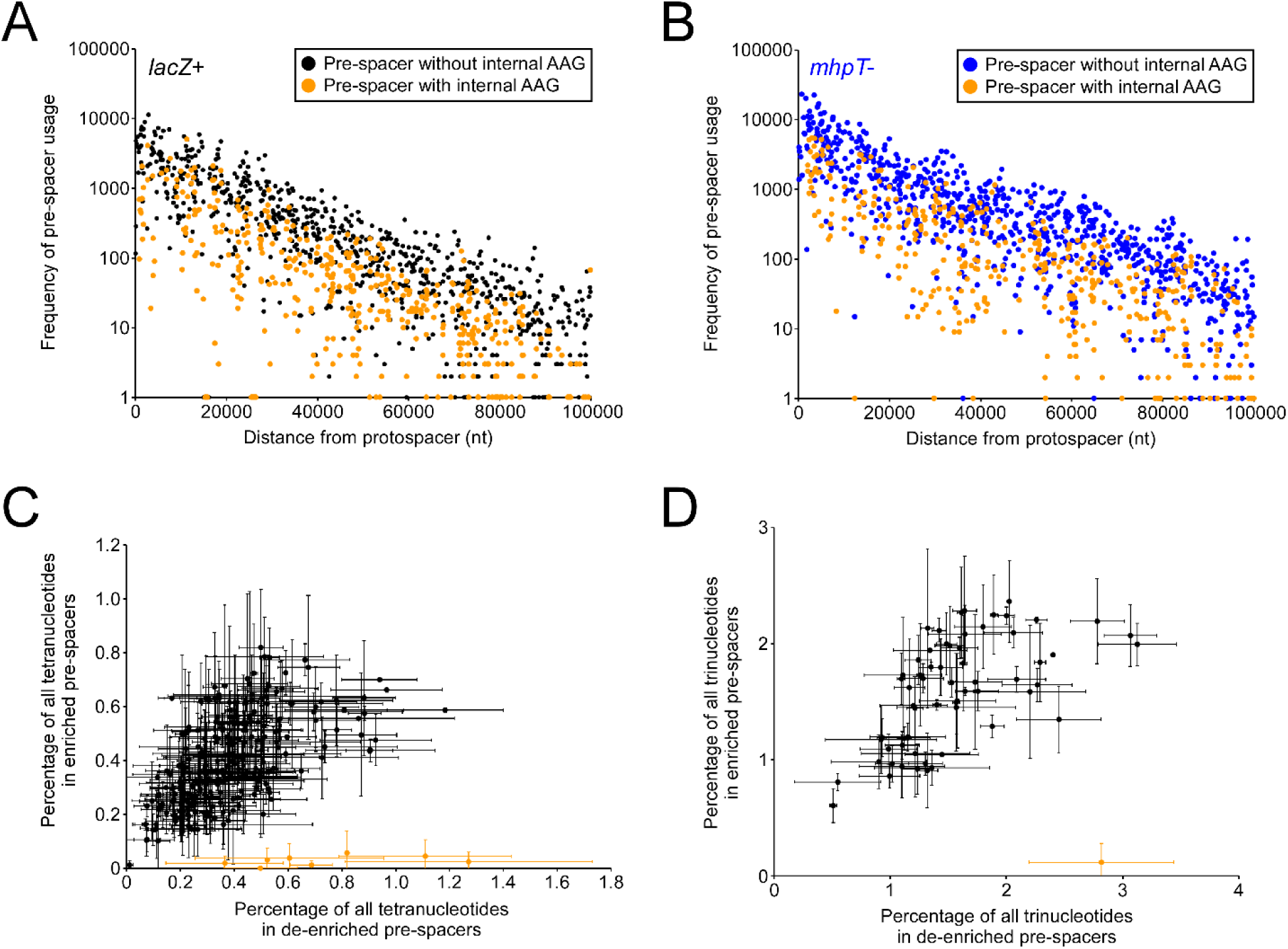
Pre-spacers that contain an AAG are used less frequently. Frequency of pre-spacer usage across the 100 kb upstream of the **(A)** *lacZ*+ and **(B)** *mhpT*-protospacers for AMD536 cells containing pAMD191 (encodes Cas3), and either pCB380 (encodes crRNA targeting *lacZ*+) or pAMD212 (encodes crRNA targeting *mhpT*-). One was added to all frequency values to allow visualization of zero-scoring datapoints. Data are only shown for 33 bp protospacers on the non-targets strand, associated with an AAG PAM. Orange datapoints represent pre-spacers with an internal AAG. **(C)** Frequency of tetranucleotide sequence usage within pre-spacers that were de-enriched (*x*-axis) or de-enriched (*y*-axis) relative to an exponential decay model. Values represent the percentage of all tetranucleotides, averaged for data from the *lacZ*+ and *mhpT*-protospacer datasets. Error bars represent one standard deviation from the mean. Tetranucleotide sequences containing AAG are plotted in orange. **(D)** Same as (C), but for trinucleotide sequences.

To experimentally test the effect of an AAG within a pre-spacer, we generated a second self-targeting plasmid, “stp2”, that includes a potential pre-spacer (AAG PAM) that can be easily modified (see Materials and Methods for a detailed description of the plasmid and assessment of its ability to cause primed adaptation). The unmodified pre-spacer in stp2 has AAA at positions 19-21. We also constructed a derivative of stp2 in which the introduced pre-spacer has been modified to contain AAG rather than AAA at positions 19-21. We refer to this mutant stp2 plasmid as stp2-mut1 (Figure 12A). We initiated primed adaptation in cells containing either stp2 or stp2-mut1. We determined the location of pre-spacers for each of the two plasmids by PCR-amplification and sequencing of the expanded CRISPR arrays. We compared the relative usage frequency for each pre-spacer on the non-target strand that is associated with an AAG PAM (these represent the most frequently used pre-spacers; Figure 2A) for each of the two plasmids. As expected, pre-spacers with the same sequence were used at similar frequencies for each plasmid (Figure 12B). By contrast, the AAG-containing pre-spacer in stp2-mut1 was used at a substantially lower frequency than the equivalent AAA-containing pre-spacer in wild-type stp2 (Figure 12B). Thus, our experimental data are consistent with the bioinformatic analysis, and show that pre-spacers with an internal AAG are used at a reduced frequency during primed adaptation. We note also that a recent, independent study reached a similar conclusion about the impact of AAG sequences within pre-spacers (Musharova et al., 2018).

**Figure 12.**
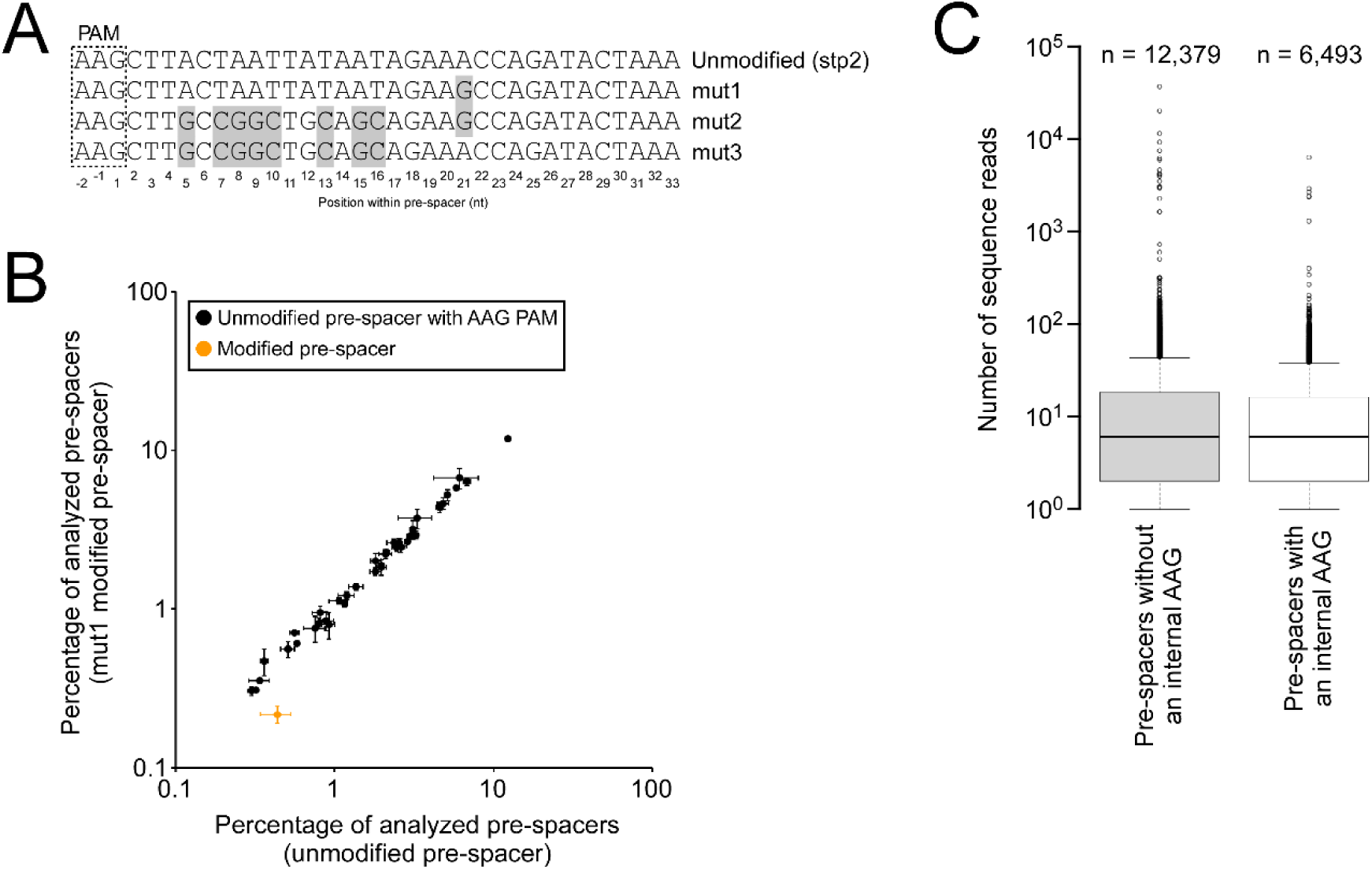
Experimental evidence that pre-spacers containing an AAG are used less frequently in primed adaptation, and to a lesser degree in naïve adaptation. **(A)** Schematic showing the unmodified and modified pre-spacer sequences for the stp2, stp2-mut1, stp2-mut2 and stp2-mut3 plasmids. The PAM is indicated by a dashed box. Grey highlighting indicates changes relative to the unmodified pre-spacer. **(B)** Frequency of pre-spacer usage for all pre-spacers on the unmodified and mut1 stp2 plasmids, for pre-spacers on the non-target strand with an AAG PAM. The single modified pre-spacer that differs between the stp2 and stp2-mut1 is shown in orange. The frequency of pre-spacer usage from the stp2 and stp2-mut1 plasmids are shown on the *x*-axis and *y*-axis, respectively. Values plotted represent the mean of two independent biological replicates. Error bars represent one standard deviation from the mean. **(C)** Tukey boxplot showing the distribution of pre-spacers usage or all chromosomal sequences with an AAG PAM that were detected at least once in an assay of naïve adaptation (Yosef et al., 2013). Pre-spacers are separated depending on the presence or absence of an internal AAG sequence.

A previous study mapped pre-spacers for naïve adaptation in a strain of *E. coli* that is not able to undergo primed adaptation or interference (Yosef et al., 2013). The authors sequenced 934,202 newly acquired spacers that came from the chromosome, covering 84,951 pre-spacer sites. Although the frequency of pre-spacers with AAG PAMs is considerably lower for naïve adaptation than primed adaptation, many pre-spacers were associated with an AAG PAM (Yosef et al., 2013). Using these data, we determined the frequency of usage for all potential chromosomal pre-spacers associated with an AAG PAM. 18,872 sequences were identified as pre-spacers at least once, whereas 108,207 sequences were not. We then compared the number of AAG-containing (i.e. an internal AAG) pre-spacers within each of these groups to determine whether the impact of an internal AAG is limited to primed adaptation. Although the proportion of AAG-containing pre-spacers was similar for each group, a significantly larger fraction (38.8%) of the unused pre-spacers had an internal AAG than the fraction of used pre-spacers (34.4%; Chi Squared Test with Yates’ Continuity Correction *p* < 1e^-7^). As a control, we analyzed the proportion of AGA-containing pre-spacers for the same groups and found no significant difference (Chi Squared Test with Yates’ Continuity Correction *p* = 0.08). We then divided the 18,872 used pre-spacer sequences into AAG-containing and non-containing groups and compared the usage frequencies for each group. While the distribution of usage frequencies was similar between the two groups (Figure 12C), it was slightly and significantly higher for the AAG-lacking group (Mann Whitney U Test *p* = 2.5e^-4^). We saw no significant difference when comparing AGA-containing and AGA-lacking pre-spacers (Mann Whitney U Test *p* = 0.63). We conclude that AAG-containing pre-spacers are acquired at a lower efficiency during naïve adaptation. However, the magnitude of the effect appears very small, in contrast to the large effect we observed for primed adaptation (Figures 11 + 12B). Consistent with this, a previous study did not detect an effect of AAG within pre-spacers on the efficiency of naïve adaptation (Musharova et al., 2018).

Since AAG is the PAM associated with >95% of pre-spacer usage during primed adaptation, we presume that the impact of AAGs within pre-spacers is connected to the selection of pre-spacers with AAG PAMs. One possible explanation for the reduced frequency of usage for AAG-containing pre-spacers is that multiple Cas3-containing complexes could translocate from a single Cascade-bound protospacer, such that stable association of a Cas3-containing complex with an AAG PAM sequence prevents association of a second Cas3-containing complex with a nearby AAG PAM. However, this would not explain the size of the decrease in pre-spacer usage frequency associated with an internal AAG (3.5-fold), or its impact over a >100 kb region; for the effect of an internal AAG to be explained by competition between Cas3-containing complexes, most of the >1,000 AAG sequences within 100 kb of the protospacer would need to be occupied at any given time. Moreover, for pre-spacers where the AAG PAM falls within an upstream pre-spacer, the frequency of usage was not significantly different to that of non-overlapping pre-spacers (Two-tailed Mann Whitney U Test *p* = 0.98 and 0.25 for *lacZ*+ and *mhpT*-, respectively), arguing against a competition model. A more likely explanation is that AAG sequences are frequently nicked, which would represent a semi-stable change in the DNA that can be caused by a transiently positioned Cas3-containing complex. Recent studies suggest that pre-spacers are nicked within the PAM during primed adaptation (Musharova et al., 2017; Shiriaeva et al., 2019). We propose that the Cas3-containing complex nicks all AAG sequences that it encounters as it translocates away from the Cascade-bound protospacer (after transitioning from the highly active state within 200 nt of the protospacer), and that pre-spacers with an internal nick cannot be used as substrates for adaptation by Cas1-Cas2.

Intriguingly, the impact of an AAG within a pre-spacer appears to diminish as the AAG is positioned further from the PAM (Figure 13A) (Musharova et al., 2018). Based on our nicking model, we hypothesized that closely positioned pairs of AAG sequences would lead to closely spaced nicks, and hence single-stranded DNA gaps that may then prevent the ability of these sequences to be integrated into a CRISPR array. To test this hypothesis, we repeated the stp2 primed adaptation assay described above (Figure 12B) using a derivative of the stp2-mut1 plasmid where the sequence between the PAM and the internal AAG of the modified pre-spacer was changed to a more G/C-rich sequence with a higher melting temperature. We reasoned that this would reduce the propensity for the DNA between the PAM and the internal AAG to become single-stranded in the event that the PAM AAG and the internal AAG were both nicked. We refer to this plasmid as stp2-mut2 (Figure 12A). We also constructed an equivalent plasmid, stp2-mut3, where the AAG within the pre-spacer was changed to an AAA (Figure 12A). Although the relative usage frequency of the modified pre-spacer was higher for stp2-mut2 than for stp2-mut1 (Figure 13B), it was also higher for stp2-mut3 than for stp2 (Figure 13C), suggesting that this effect is independent of the internal AAG, and arguing against our hypothesis. Moreover, these data suggest that the sequence of the PAM-proximal region of a pre-spacer impacts the frequency with which it is used during primed adaptation.

**Figure 13.**
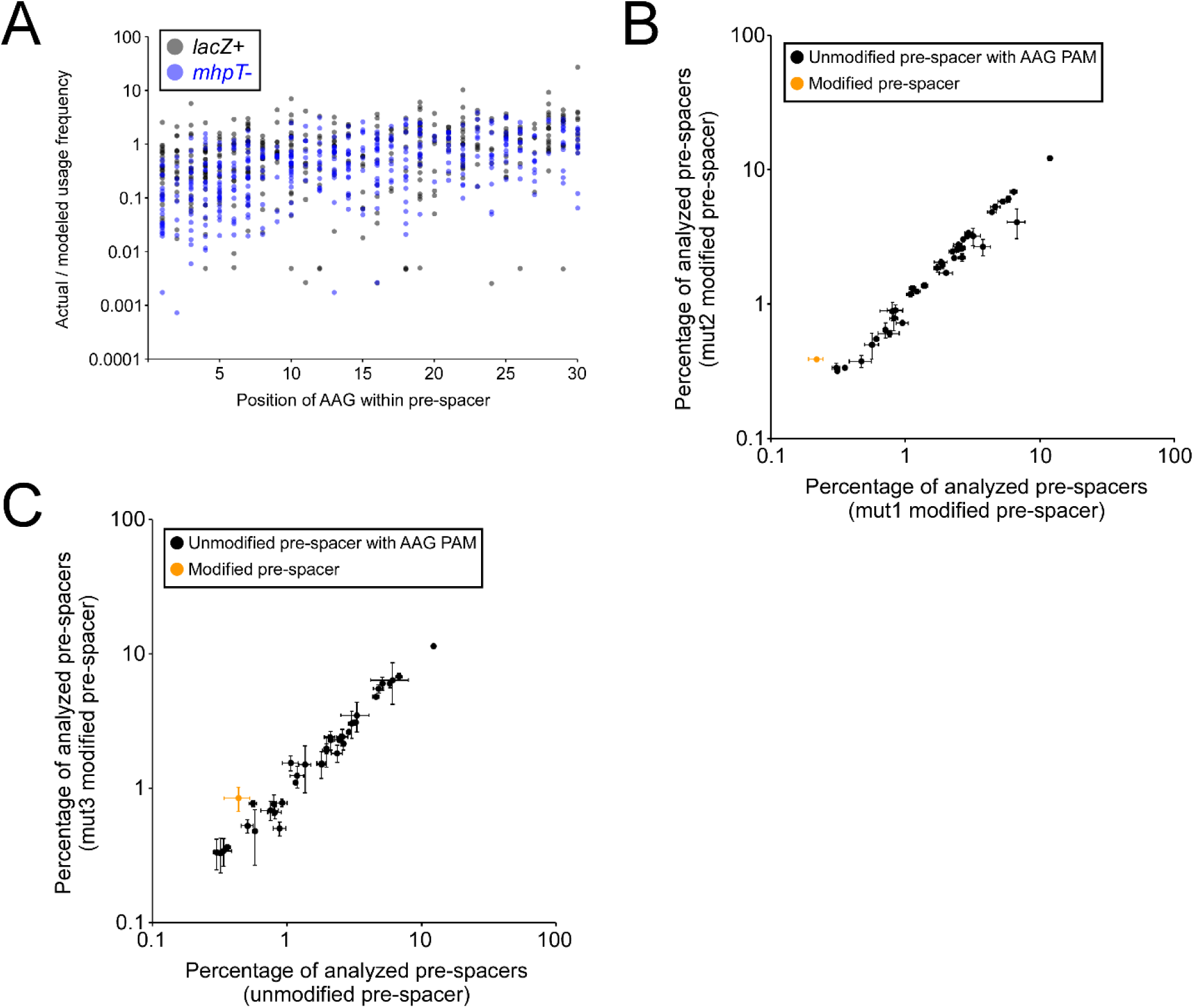
The impact of an AAG within a pre-spacer on primed adaptation efficiency is dependent on position of the AAG relative to the PAM. **(A)** Ratio of actual pre-spacer usage frequency to the frequency predicted by an exponential decay model for all pre-spacers with an internal AAG. Data are only shown for pre-spacers with an AAG PAM, on the non-target strand, <100 kb upstream of the *lacZ*+ (black datapoints) or *mhp*T- (blue datapoints) protospacer. Ratios are shown as a function of the position of the internal AAG sequence within the pre-spacer. **(B)** Frequency of pre-spacer usage for all pre-spacers on the stp2-mut1 and stp2-mut2 plasmids, for pre-spacers on the non-target strand with an AAG PAM. The single modified pre-spacer that differs between stp2-mut1 and stp2-mut2 is shown in orange. The frequency of pre-spacer usage from the stp2-mut1 and stp2-mut2 plasmids are shown on the *x*-axis and *y*-axis, respectively. **(C)** As for (B) but for the stp2 and stp2-mut3 plasmids. For (B) and (C), values plotted represent the mean of two independent biological replicates. Error bars represent one standard deviation from the mean.

### The efficiency of primed adaptation is modulated by a PAM-distal motif in the pre-spacer

Although the presence of an AAG within the pre-spacer is clearly an important factor in determining the frequency of pre-spacer usage, there is considerable local variability in pre-spacer usage for pre-spacers that lack an internal AAG (Figure 11A + B). We used the frequency of pre-spacer usage for AAG-flanked pre-spacers lacking an internal AAG, in the 100 kb region upstream of the *lacZ*+ and *mhpT*-protospacers, to fit exponential decay models, estimating pre-spacer usage as a function of distance from the protospacer (Formulae of exponential decay models for *lacZ*+ and *mhpT*-, respectively, are *y* = 1541.8e^-6E-05*x*^ and *y* = 4073.7e^-5E-05*x*^, where *y* is the pre-spacer usage frequency and *x* is the distance from the protospacer; R^2^ values for the *lacZ*+ and *mhpT*-models, respectively, are 0.51 and 0.66). Based on the modeled exponential decay, we estimate that the frequency of pre-spacer usage drops by 50% every ∼11,500 – 13,900 kb. Strikingly, an *in vitro* study of *E. coli* Cas3 translocation indicated that 50% of Cas3 molecules dissociate from DNA after ∼12 kb (Redding et al., 2015), strongly suggesting that the decrease in pre-spacer usage we observe as a function of distance from the protospacer reflects the extent of Cas3 translocation.

We compared the frequency of pre-spacer usage to the predicted frequency based on the exponential decay model. We then selected all pre-spacers for which the frequency of usage was >4-fold greater than that predicted by the model. Alignment of these pre-spacer sequences revealed sequence bias at positions 28, 30, 32 and 33 (Figure 14A). We next selected all pre-spacers for which the frequency of usage was >4-fold lower than that predicted by the model. Alignment of these pre-spacer sequences revealed sequence bias at positions 30, 31, 32 and 33 (Figure 14B). Strikingly, the hierarchy of preferred bases at positions 30, 32 and 33 was precisely inverted for the enriched and de-enriched pre-spacers. To determine whether the identified sequence biases are meaningful, we determined the information content of the position weight matrices (Schneider et al., 1986) for the enriched and de-enriched pre-spacers. We then determined the information content of the same number of randomly selected pre-spacer sequences, and repeated this analysis 100,000 times. Based on the distribution of information content scores for the randomly selected sequences, the information content scores of the position weight matrices from the enriched and de-enriched pre-spacers were significantly higher than expected by chance (*Z* = 10.4 and *p* < 1e^-14^ for enriched pre-spacers; *Z* = 4.9 and *p* = 5e^-7^ for de-enriched pre-spacers). We conclude that the sequence of base positions at the 3’ end of the pre-spacer influences usage frequency, with positions 30, 32 and 33 being the most important.

**Figure 14.**
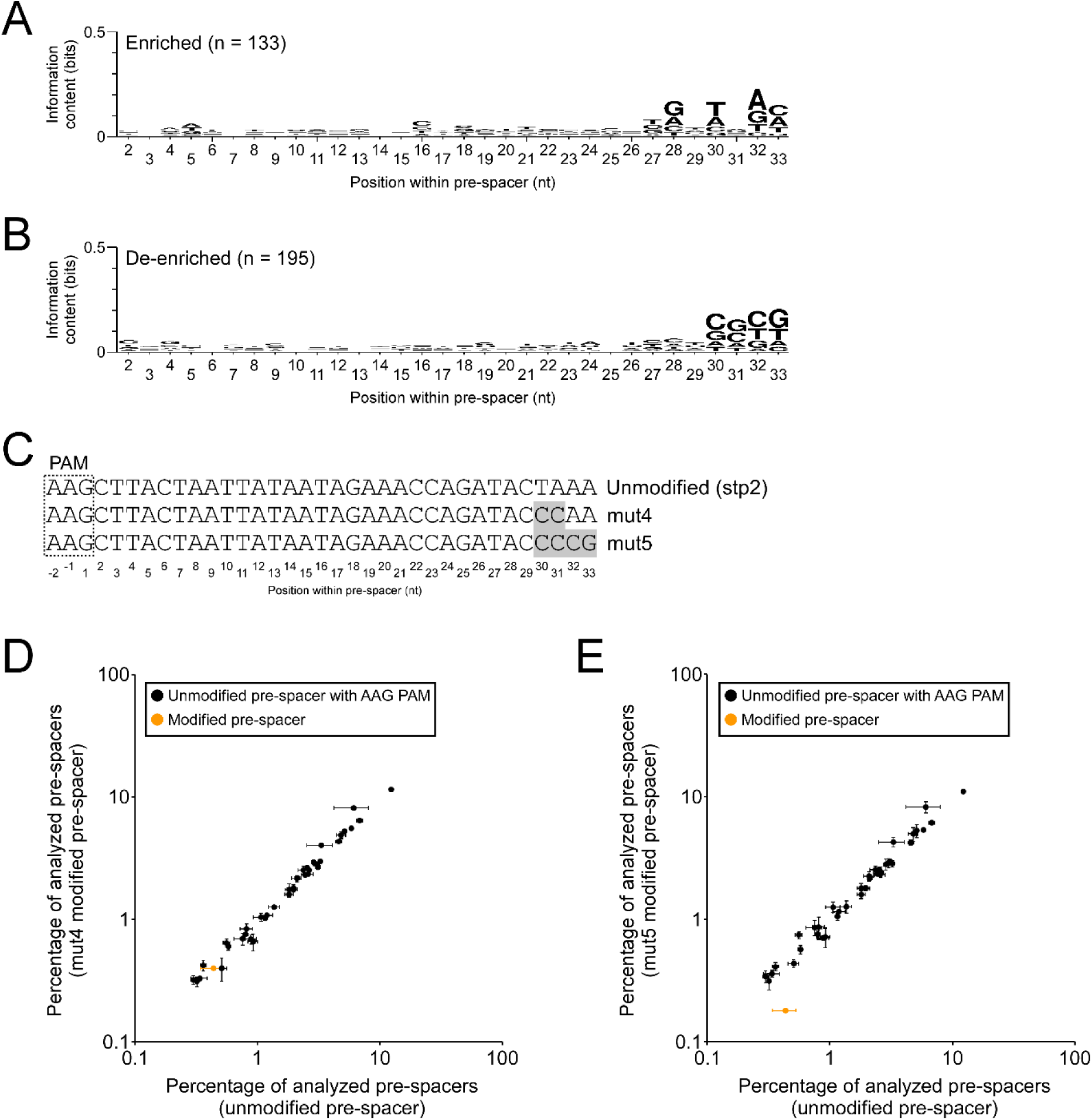
Sequences at the PAM-distal end of the pre-spacer impact the efficiency of primed adaptation. **(A)** DNA sequence logo for the 133 “enriched” pre-spacers without an internal AAG that are used >4-fold more frequently than expected based on the exponential decay model. Note that the *y*-axis maximum is set to 0.5 bits. **(B)** DNA sequence logo for the 195 “de-enriched” pre-spacers without an internal AAG that are used >4-fold less frequently than expected based on the exponential decay model. Note that the *y*-axis maximum is set to 0.5 bits. **(C)** Schematic showing the unmodified and modified pre-spacer sequences for the stp2, stp2-mut4, and stp2-mut5 plasmids. The PAM is indicated by a dashed box. Grey highlighting indicates changes relative to the unmodified pre-spacer. **(D)** Frequency of pre-spacer usage for all pre-spacers on the stp2 and stp2-mut3 plasmids, for pre-spacers on the non-target strand with an AAG PAM. The single modified pre-spacer that differs between stp2 and stp2-mut4 is shown in orange. The frequency of pre-spacer usage from the stp2 and stp2-mut4 plasmids are shown on the *x*-axis and *y*-axis, respectively. **(E)** As for **(D)** but for the stp2 and stp2-mut5 plasmids. For (D) and (E), values plotted represent the mean of two independent biological replicates. Error bars represent one standard deviation from the mean.

To experimentally test the impact of pre-spacer positions 30-33 on primed adaptation, we constructed derivatives of stp2 that contain a pre-spacer where positions 30-31 were modified from TAAA to CCAA (stp2-mut4), or positions 30-33 were modified to CCCG (stp2-mut5; Figure 14C). Based on the enrichment/de-enrichment of pre-spacers in the chromosomal primed adaptation experiment (Figure 14A-B), we reasoned that changing positions 30-33 to CCAA would not have a large effect on the frequency of pre-spacer usage, since our data suggest that only one of the two modified bases (position 30) would generate a sub-optimal pre-spacer. By contrast, we reasoned that changing positions 30-33 to CCCG would have a substantial negative impact on the frequency of pre-spacer usage, since it has the most disfavored base at three PAM-distal positions. Consistent with our expectation, changing positions 30-31 to CC had no detectable effect on pre-spacer usage frequency (Figure 14D), whereas changing positions 30-33 to CCCG substantially reduced pre-spacer usage frequency (Figure 14E).

A previous study suggested that the sequence of positions 32 and 33 of the pre-spacer influences the efficiency of naïve adaptation. Specifically, a C at position 32 or T at position 33 resulted in inefficient adaptation whereas an A at position 32 and or an A at position 33 resulted in efficient adaptation (Yosef et al., 2013). Our data are consistent with the idea that the sequence of positions 28-33 are important, and moreover are consistent with the specific sequence effects observed for naïve adaptation. We conclude that the sequence of the PAM-distal portion of the pre-spacer influences both naïve and primed adaptation, likely due to effects on Cas1-Cas2 association with, and/or integration of, the pre-spacer into the CRISPR array. Our data suggest that the number of important positions at the PAM-distal end of the pre-spacer is larger than previously suggested (Yosef et al., 2013), encompassing perhaps as many as six positions.

### A model for primed adaptation

We propose that once recruited to a protospacer-bound Cascade, Cas3 translocates along the DNA, upstream from the PAM, consistent with *in vitro* studies (Dillard et al., 2018; Redding et al., 2015). The initial translocating state of Cas3 is likely to be different to its final translocating state; we propose that this initial state is associated with frequent, sequence-independent nicking of one strand of the DNA. Our data suggest that the transition from the frequently nicking state to the less frequently nicking state is stochastic, and occurs in all cases within ∼200 nt of the protospacer. Biochemical data suggest that translocating Cas3 initially remains associated with Cascade, reeling in DNA rather than moving along it freely (Dillard et al., 2018; Loeff et al., 2018; Redding et al., 2015). We propose that the tethering of Cas3 to Cascade causes Cas3 to be in the frequently nicking state, and that it is the dissociation of Cas3 from Cascade as tension builds in the DNA that causes Cas3 to transition to the less frequently nicking state. After the transition, we propose that Cas3, or possibly Cas1 that is associated with Cas3, nicks DNA at AAG/CTT sequences. Given the magnitude of the defect in primed adaptation associated with the presence of an AAG within a spacer, it is likely that this nicking occurs at almost every AAG/CTT encountered by the Cas3-containing complex. It is unclear why the degree to which an AAG within a pre-spacer affects primed adaptation is dependent on the position of the AAG relative to the PAM. Given that an AAG within a pre-spacer has a small but significant negative impact on naïve adaptation, we suggest that Cas1 is responsible for the DNA cut at CTT sequences (Cas1 recognizes the CTT-containing strand rather than the AAG). Presumably, during naïve adaptation, a second cut by Cas1 is less likely than during primed adaptation, because Cas1 would not be associated with a translocating Cas3, and hence is less likely to be suitably positioned on the DNA.

It is unclear how substrates are generated for Cas1-Cas2 during primed adaptation, although nicking of at least one DNA strand by Cas3 is required (Datsenko et al., 2012). There is strong evidence for a nick between the second and third positions of the PAM, on the AAG-containing strand (Musharova et al., 2017; Shiriaeva et al., 2019), consistent with our proposal that every AAG is cut by the translocating Cas3-containing complex. However, this would only account for one of the four cuts required for substrate generation. Future work is required to determine whether Cas3, Cas1, and/or non-Cas proteins are required for this process. Once substrates have been generated, the efficiency of integration into the array is dependent on the sequence. Our data indicate that the six most PAM-distal positions of the pre-spacer contribute to the efficiency of primed adaptation (Figure 14). Given that the importance of the two most PAM-distal positions has been recognized to be important for naïve adaptation (Yosef et al., 2013), we propose that the impact of these positions is solely on Cas1-Cas2. These positions may be important for stable association of Cas1-Cas2, or for the process of integration into the CRISPR array. Our data also suggest that sequences within pre-spacers, more proximal to the PAM, play an important role in primed adaptation (Figure 13), but the mechanism for this is unknown.

## MATERIALS AND METHODS

### Strains and Plasmids

Strains, plasmids, and oligonucleotides are listed in Tables 1, 2, and S1, respectively. All strains are derivatives of MG1655 (Blattner et al., 1997). CB386, AMD536 and AMD688 have been previously described (Cooper et al., 2018; Luo et al., 2014). 1XDNAi, as described previously (Caliando and Voigt, 2015), contains a chromosomally integrated actuator, pACT-01. AMD671 is a derivative of 1XDNAi that we constructed using FRUIT recombineering (Stringer et al., 2012). Specifically, *thyA* was deleted in 1XDNAi, using oligonucleotides JW472 and JW473 and strain AMD052 (MG1655 Δ*thyA*) as a template. *thyA* was inserted downstream of the pACT-01 actuator using FRUIT (Stringer et al., 2012) with oligonucleotides JW9016 and JW9017. Oligonucleotides JW9009 and JW9010 were used to amplify *cas1*-*cas2* from MG1655 to replace *thyA* downstream of pACT-01. Lastly, the *lacZ* gene was replaced by *thyA* using FRUIT (Stringer et al., 2012) with oligonucleotides JW9066 and JW9067.

**Table 1.** Strains used in this study.

| Name | Description | Source |
| --- | --- | --- |
| MG1655 | Wild-type *E. coli* | (Blattner et al., 1997) |
| CB386 | MG1655 [Δ*cas3* *PcseI*]::[*cat* PJ23119] | (Luo et al., 2014) |
| AMD536 | MG1655 [Δ*cas3* *PcseI*]::[ PJ23119] | (Cooper et al., 2018) |
| AMD688 | MLS1003 [Δ*cas3* *PcseI*]::[ PJ23119] | (Cooper et al., 2018) |
| 1XDNAi | MG1655 Δ*araC-araBAD*, Δ*PlacI:lacI*, Δ*cas-CRISPR::pACT-01* (*kan*<sup>r</sup>) | (Caliando and Voigt, 2015) |
| AMD671 | MG1655 Δ*araC-araBAD*, Δ*PlacI:lacI*, Δ*cas-CRISPR::pACT-01*, *cas1*<sup>+</sup> *cas2*<sup>+</sup>, Δ*lacZ::thyA* | This Study |
| AMD052 | MG1655 Δ*thyA* | (Stringer et al., 2012) |

**Table 2.** Plasmids used in this study.

| Name | Description | Source |
| --- | --- | --- |
| pBAD24 amp | Empty pBAD24 amp | (Guzman et al., 1995) |
| pBAD33 cam | Empty pBAD33 cam | (Guzman et al., 1995) |
| pAMD179 | Parent vector for cloning crRNAs | (Cooper et al., 2018) |
| pAMD189 | stp; pAMD179 expressing a self-targeting crRNA | (Cooper et al., 2018) |
| pAMD191 | pBAD33-*cas3* | (Cooper et al., 2018) |
| pCB380 | pcrRNA.con-*lacZ* | (Luo et al., 2014) |
| pAMD211 | pAMD179 expressing a crRNA targeting *mhpT* | This Study |
| pAMD212 | pAMD179 expressing a crRNA targeting *codA* | This Study |
| pSDS009 | Parent vector for cloning pre-spacers | This Study |
| pSK013 | pSDS009 stp2-mut1 pre-spacer | This Study |
| pSK014 | stp2; pAMD179 with an added pre-spacer | This Study |
| pSK015 | pSDS009 stp2-mut2 pre-spacer | This Study |
| pSK035 | pSDS009 stp2-mut3 pre-spacer | This Study |
| pSK016 | pSDS009 stp2-mut5 pre-spacer | This Study |
| pSK017 | pSDS009 stp2-mut4 pre-spacer | This Study |

The self-targeting plasmid (stp; pAMD189), the Cas3-expressing plasmid (pAMD191), and the plasmid that expresses the crRNA targeting the *lacZ*+ protospacer (pCB380), have been described previously (Cooper et al., 2018; Luo et al., 2014). All other crRNA-expressing plasmids are derivatives of pAMD179 (Cooper et al., 2018). To clone individual spacers, pairs of oligonucleotides were annealed, extended, and cloned using In-Fusion (Clontech) into the *<u>XhoI and SacII</u>* sites of pAMD179 to generate pAMD211 (targets *codA*+ protospacer; with oligonucleotides JW8010 and JW6518), and pAMD212 (targets *mhpT*-protospacer; with oligonucleotides JW8011 and JW6518).

The second self-targeting plasmid (stp2) is a derivative of pSDS009. pSDS009 was generated by cloning a synthesized dsDNA fragment (gBlock JW9076; Integrated DNA Technologies, Inc.; Table S1) using In-Fusion (Clontech) into the *Cla*I and *Hin*dIII sites of pBAD24 amp (Guzman et al., 1995). stp2 and its derivatives were generated by annealing and extending pairs of oligonucleotides, and cloning using In-Fusion (Clontech) into the *<u>Hin</u>*<u>dIII</u> site of pSDS009 to generate pSK014 (stp2; cloned with oligonucleotides JW10011 and JW10012), pSK013 (stp2-mut1; cloned with oligonucleotides JW10009 and JW10010), pSK015 (stp2-mut2; cloned with oligonucleotides JW10013 and JW10014), pSK035 (stp2-mut3; cloned with oligonucleotides JW10015 and JW10016), pSK017 (stp2-mut4; cloned with oligonucleotides JW10021 and JW10022), and pSK016 (stp2-mut5; cloned with oligonucleotides JW10017 and JW10018).

### Quantification of adaptation using a YFP reporter

AMD688 was transformed with two plasmids: the first plasmid was either pAMD189 or empty pBAD24, while the second was either pAMD191 or empty pBAD33. Cells were grown overnight in LB supplemented with 100 ug/mL ampicillin and 30 ug/mL chloramphenicol and were sub-cultured the next day 1:100 for 6 hours in LB supplemented with 0.2% arabinose and 30 ug/mL chloramphenicol. Cells were pelleted by centrifugation and resuspended in M9 minimal medium in twice the original volume (OD_600_ ≈ 1.0). Samples were transferred to 5 mL polystyrene round-bottom tubes and analyzed by flow cytometry for single-cell detection of YFP expression using the BD FACSAria IIU Cell Sorter. To remove debris and aggregated cells, we gated cells on (i) Forward Scatter Area and Side Scatter Area, (ii) Side Scatter Height and Side Scatter Width, and (iii) Forward Scatter Height and Forward Scatter Width. We then measured fluorescence of the remaining cells. We recorded 100,000 events for each sample, of which 70-90% were retained after the gating steps (Supplementary Dataset).

### PCR and Sequencing to Assess Primed Adaptation

To determine the set of pre-spacers acquired during priming, pAMD191 was transformed into AMD536 along with either pAMD189, pCB380, pAMD211 or pAMD212. Cells were grown overnight in LB supplemented with 100 ug/mL ampicillin, 30 ug/mL chloramphenicol, and 0.2% glucose at 37 °C with aeration, and sub-cultured the next day in LB supplemented with 0.2% arabinose at 37 °C with aeration for one hour.

To experimentally assess the impact of specific sequences on pre-spacer usage, pSK013, pSK014, pSK015, pSK035, pSK016, or pSK017 were transformed into AMD671. Cells were grown overnight in LB supplemented with 100 ug/mL ampicillin, and 0.2% glucose at 37°C with aeration, and sub-cultured the next day in LB supplemented with 0.2% arabinose at 37°C with aeration for six hours.

For both sets of cultures described above, cells were pelleted from 1 mL of culture by centrifugation, and cell pellets were frozen at -20 °C. PCRs were then performed on the cell pellets, amplifying the CRISPR arrays using oligonucleotides JW7816 and JW7817 for CRISPR-I and JW7818 and JW7819 for CRISPR-II.

The amplicon sizes for unexpanded CRISPR-I and CRISPR-II arrays are 237 bp and 236 bp, respectively. PCR products were visualized on acrylamide gels and the first expansion band was extracted and purified. A second round of amplification was performed using a universal forward oligonucleotide (JW7820 or JW8053), and one of a set of reverse oligonucleotides containing Nextera (Illumina) indices (JW8054, JW8057, JW8062, JW8476, JW8477, JW8478, JW8479, JW8480, JW8481 or JW8485). Samples were purified using a MinElute kit (Qiagen) and pooled. Samples were sequenced using an Illumina MiSeq Instrument (Wadsworth Center Applied Genomic Technologies Core). Sequence reads were 250 nt.

Plasmids pSK013, pSK014, pSK015, pSK035, pSK016, or pSK017 serendipitously function as self-targeting plasmids despite the lack of a cloned spacer. These plasmids are derivatives of pSDS009, which is designed to express crRNAs from a cassette that includes a spacer with a full repeat downstream, but only 8 bp of repeat sequence upstream. pSDS009 does not itself contain a canonical spacer between the partial and complete repeat sequences. Similarly, pSK013, pSK014, pSK015, pSK035, pSK016, and pSK017 do not include a canonical spacer sequence. Nonetheless, multiple lines of evidence indicate that these plasmids drive efficient primed adaptation in cells expressing *cas3*: (i) the majority of pre-spacer usage is from the plasmid; (ii) 96% of pre-spacer usage on the plasmid is associated with an AAG PAM; (iii) for AAG-flanked pre-spacers that are shared with the stp, are found on the stp non-target strand, and are not shared with pAMD191, pre-spacer usage from pSK014 correlates well with that from the stp (correlation coefficient of 0.93). We suspect that the downstream repeat sequence in pSDS009 and its derivatives also functions as an upstream repeat sequence for a cryptic crRNA. In this scenario, the sequence downstream of this repeat serves as a spacer, with the end of the crRNA presumably being determined by the position of the downstream transcription terminator. We previously showed that the exact same sequence results in expression of a cryptic crRNA from a derivative of pAMD179 (Cooper et al., 2018). The presumed spacer sequence is duplicated in the same orientation, a short distance downstream, which would facilitate interference and primed adaptation. The presumed protospacer has an AGT PAM.

### Spacer sequence extraction and alignment to reference genomes

Newly acquired spacers found directly adjacent to the leader proximal endogenous spacer (the first primed adaptation acquired spacer) were extracted from .fastq files using custom Python scripts for each CRISPR array being analyzed. The scripts first selected sequences that included the first spacer of the unexpanded CRISPR array (spacer #1) and at least one repeat sequence. The script then removed sequences for which the gap between spacer #1 and the next upstream repeat differed from the expected value. The script then extracted the sequence between the first repeat and the repeat immediately upstream of spacer #1. This was presumed to consist of complete spacer sequences, or combinations of spacers and repeats (i.e. the result of >1 spacer being added to the CRISPR array). Sequences were discarded if they were shorter than 30 nt or longer than 36 nt, ensuring that only arrays with a single expansion were analyzed. Spacers were mapped to their respective genomes (stp, pAMD191, or *E. coli* K-12 MG1655 U00096.3) using CLC Genomics Workbench v10.1.1, requiring perfect matches. For experiments involving mapping reads to multiple genomes, we first mapped to stp, then mapped all unmapped reads to pAMD191, and then mapped all unmapped reads to *E. coli* K-12 MG1655.

### Analysis of primed adaptation in regions adjacent to off-target Cascade binding sites

We previously described a set of 76 off-target Cascade binding sites whose sequence is consistent with Cascade binding in association with the crRNA targeting the *lacZ*+ protospacer (Cooper et al., 2018). We counted the frequency of pre-spacer usage for each of these off-target protospacers, searching on the non-target strand in the 10 kb upstream of the protospacer. We repeated the analysis for 1,000 randomly selected genomic positions equally distributed across the forward and reverse strands.

### Analysis of slipping for 33 bp pre-spacers

We selected all 33 bp pre-spacers on the non-target strand, in the 10 kb regions upstream of the *lacZ*+ and *mhpT*-protospacers, for which the PAM was not AAG. We calculated all pairwise distances for these pre-spacers to AAG-associated pre-spacers on the non-target strand, in the 10 kb upstream of each of the protospacer. For the analysis represented in Figure 7A, only unique pre-spacer positions were considered, so information on the frequency with which different pre-spacers are used was lot. Values plotted in Figure 7A represent the number of instances of the indicated slipping distance divided by the total number of comparisons between pre-spacer positions and AAGs, multiplied by 1000. Values listed in Figure 7B take into account the fact that many pre-spacer positions were detected more than once, and hence these numbers control for the relative use of the different sequences. Pre-spacers with a non-AAG PAM that did not derive from a -1, +1 or +2 slip were selected; pre-spacers that were sequenced multiple times were represented at the frequency with which they were detected. The associated PAM sequences were then converted into the logos shown in Figure 7C using Weblogo (Crooks et al., 2004).

### Analysis of flipping

We selected all 33 bp pre-spacers on the target strand, in the 10 kb regions downstream of the *lacZ*+ and *mhpT*-protospacers. We calculated all pairwise distances for these pre-spacers to AAG-associated pre-spacers on the non-target strand, in the 10 kb upstream of each of the protospacer. For the analysis represented in Figure 8B, only unique pre-spacer positions were considered, so information on the frequency with which different pre-spacers are used was lot. Values plotted in Figure 8B represent the number of instances of the indicated slipping distance divided by the total number of comparisons between pre-spacer positions and AAGs, multiplied by 1000.

### Analysis of slipping for spacers of non-canonical length

We selected all 32 and 34 bp pre-spacers on the non-target strand, in the 10 kb regions upstream of the *lacZ*+ and *mhpT*-protospacers. We calculated all pairwise distances for these pre-spacers to AAG-associated pre-spacers on the non-target strand, in the 10 kb upstream of each of the protospacer. For the analysis represented in Figure 9C-D, only unique pre-spacer positions were considered, so information on the frequency with which different pre-spacers are used was lot. Values plotted in Figure 9C-D represent the number of instances of the indicated slipping distance divided by the total number of comparisons between pre-spacer positions and AAGs, multiplied by 1000.

### Identification of pre-spacers with higher-than-expected or lower-than-expected usage frequency, independent of internal AAG sequences

We took the frequency of pre-spacer usage for all AAG-flanked 33 bp pre-spacers in the 100 kb region upstream of the *lacZ*+ and *mhpT*-protospacers. We then fit an exponential decay model to each dataset using Microsoft Excel (“best fit line” function). Using the parameters associated with these models, we determined the expected usage frequency for pre-spacers at every position in the 100 kb windows, and we then took the ratio of the actual usage frequency to the expected usage frequency. Pre-spacers with ratios >4 or <0.25 were selected for analysis of tetranucleotide and trinucleotide content.

### Identification of pre-spacers with higher-than-expected or lower-than-expected usage frequency, independent of internal AAG sequences

We determined expected pre-spacer usage frequency by fitting an exponential decay model, as described above, but first excluding pre-spacers with an internal AAG. We then took the ratio of the actual usage frequency to the expected usage frequency. Pre-spacers with ratios >4 or <0.25 were used as input for Weblogo (Crooks et al., 2004) to create the sequence logos shown in Figure 14A-B.

## Supporting information

Reviews with point-by-point response

Marked-up copy showing changes to the original submission in response to the reviews

## ACKNOWLEDGEMENTS

We thank the Wadsworth Center Applied Genomic Technologies Core Facility for DNA sequencing. We thank the Wadsworth Center Immunology Core Facility for flow cytometry analysis. We thank the Wadsworth Center Tissue Culture and Media Core Facility and Glassware Facility for technical support. We thank Randy Morse, Keith Derbyshire and Todd Gray for helpful discussions.

## ACCESSION NUMBER

Raw Illumina sequencing data are available from EBI ArrayExpress using accession number E-MTAB-8700.

