## Supplementary material for "Characterization of Primed Adaptation in the *Escherichia coli* type I-E CRISPR-Cas System": Reviews with point-by-point response

This document includes three reviews from expert reviewers selected by Ailong Ke (Cornell University), and our response to the reviews. Text from the reviews is in black. Our responses are in red. All changes made to the original submission in response to the reviews are indicated in red text in an unformatted copy of the manuscript that is included as a supplementary file.

**Reviewer #1:**

Stringer et al. have posted an interesting manuscript describing an extremely detailed analysis of the types of spacers acquired through primed acquisition in the type I-E CRISPR-Cas system of *E. coli*. Primed adaptation was discovered in 2012 and has been the subject of considerable study in the meantime. However, most previous priming studies were performed against plasmid DNA. Because plasmids are short, there are a limited number of spacers that can be acquired and limited amount of analysis that can be done on location and sequence-specific effects on spacer acquisition. To overcome these issues, the authors performed self-targeting priming experiments, in which multiple crRNAs targeting "priming" protospacers (e.g. protospacers with a priming PAM) in the *E. coli* genome were used for priming assays followed by high-throughput sequencing (HTS) of the resulting acquired spacers. These experiments show significant primed adaptation against the genomic DNA. The authors performed extensive analysis of these spacers, focusing on the location from which they were derived, the types of PAMs that are associated with the spacers, the effect of PAM "slipping" during spacer acquisition, and the effect of the prespacer sequence on the frequency of acquisition of that sequence.

The analysis in this paper is generally rigorous and the experiments are well-performed. The results recapitulate results from previous studies, and in some cases provide new interesting insight (especially the data in Figure 14, see major concern 10 below). Overall, this work serves the important role of reproducing results from plasmid studies, while also revealing some new insights from the more information-rich source of spacers. We have several comments on data analysis and presentation that we hope the authors will be able to address in a revised manuscript.

Major concerns

1. Shiriaeva et al. 2019 also performed similar experiments using a self-targeting crRNA to analyze spacer acquisition through self-priming. They observed much stronger bidirectional spacer acquisition on opposite strands than is shown in Fig. 3 and 6a of the current manuscript (Fig. 1e of that paper). One difference between the two studies is that the crRNA in the previous study targeted a region with a correct PAM, while this study targeted regions with "priming" PAMs. This discrepancy could suggest a mechanistic difference in how prespacers are generated (or how Cas3 is recruited/activated) that depends on the PAM sequence.

We see a difference in the ratio of spacers acquired from the target vs non-target strand not only between our data and those of Shiriaeva *et al.*, but also between our data for the different protospacers (see Figure 3 and Figure 6B). We do not know the cause of these differences. We don't think it's as simple as the PAM. Shiriaeva *et al.* targeted a protospacer with an AAG PAM but a mismatch in the first position of the protospacer relative to the spacer. All three chromosomal protospacers in our study have an AGG PAM and a perfect match to the corresponding spacer. It's not straightforward to compare the effects of mismatches in the seed sequence to alterations to the PAM, since these can be protospacer-dependent (Xue *et al.*, 2015). Given the differences we see between the different protospacers in our own study, all of which use the same PAM, we suspect the effect is not due to the PAM. Whatever causes these differences is clearly important in the mechanism of priming; hence, we have expanded our discussion of this phenomenon to mention the work of Shiriaeva *et al*.

2. Related to the above comment, have the authors considered the consequences of self-spacer acquisition on selective pressure in their system? In a strain with a fully-functioning CRISPR-Cas system, acquisition of self-spacers should cause autoimmunity. Indeed, in Shiriaeva et al. 2019, the authors observed a deletion of the chromosomal region containing the self-targeted protospacer and pronounced growth defects. It is also possible that some spacers cause more autoimmunity than others (e.g. spacers acquired from an essential gene or that promote more efficient interference). This could impact which spacers are more abundant in the population. For example, the analysis shown in Figure 4 may also need to consider whether any of the Cascade off-target binding sites were in essential regions of the genome. If that were the case, it is possible that priming in these regions would face strong negative selection. Similarly, it is possible that an internal AAG sequence enhances interference, so spacers acquired from regions with internal AAGs may be negatively

selected against. The authors should include some discussion of the effects of negative selection depending on the type of spacer that is acquired during self-priming, and how this may affect their data/analysis.

This is an important point that we now address in the text. In data that we haven't included in this manuscript, we also see local deletions at chromosomal protospacers under conditions where interference occurs, presumably because of selection for deletion mutants due to the lethality of double-strand breaks caused by interference. We expect protospacers anywhere on the chromosome to be lethal, but if a protospacer is in an essential gene, the lethality of interference cannot be prevented by a local deletion. The distribution of pre-spacer usage as a function of distance from the protospacer (Figure 11A+B) does not show any obvious decreases in gene-sized regions. To be more rigorous, we looked in the *secD*, *secF*, *ribD*, *ribE* and *thiL* genes, which are all essential and are all in the "priming zone" of the *lacZ*+ protospacer. There is no detectable decrease in the efficiency of pre-spacer usage within these genes relative to the surrounding, non-essential genes ($p$ = 0.55). With respect to the off-target protospacers in Figure 4, very few of these are in essential genes. Lastly, our priming data using an AAG-containing pre-spacer in a plasmid that is not under antibiotic selection (Figure 12) are consistent with the conclusion that AAG sequences within pre-spacers directly impact priming efficiency.

3. The YFP reporter assay used in Figure 1 is highly sensitive for detecting cells that have acquired spacers. However, it seems that the overall rate of priming is much lower for this strain than those reported in other studies where spacer acquisition was measured in the native CRISPR array, including the authors' previous work (for example, Fig. 4B of Cooper et al. 2018). Did the authors observe similarly low levels of priming into the native CRISPR arrays when using the stp for the experiment shown in Figure 2? Have the authors tested expansion of the minimal CRISPR in the yfp strain by PCR for the overall population? It would be helpful if the authors could provide an example of the PCR for each type of experiment (i.e. into the minimal and native CRISPRs), if they have that data readily available.

We have measured spacer acquisition in both native CRISPR arrays for cells targeting the stp (see supplementary Figure S1 of Cooper *et al.*) in cells lacking the *yfp* cassette, and it is much more efficient than acquisition into the *yfp* locus. We have not determined why this is, but we suspect that competition with the native CRISPR-I array (present in the strain used for the experiments described in Figure 1) plays a part. We now discuss this in the text.

4. Related to the above comment, it would be helpful to show some flow cytometry data for the YFP+ cells in Figure 1. PCR to show CRISPR expansion of only the YFP+ cells would be a good addition (if the data is available), to show that the YFP+ cells did indeed all contain a newly acquired spacer.

We have included a supplementary file with the output files for all the flow cytometry experiments. We did not PCR from the cells because we had a technical problem when trying to collect the sorted cells.

5. Related to the above comments, differences have been observed in priming between studies in which *cas* genes are overexpressed versus, for example, low-level constitutive expression in *hns* deletion strains. The authors use a strain in which all *cas* genes except for *cas3* were expressed from a relatively strong constitutive promoter (J23119 from Parts Registry). It is possible that the expression levels of these genes affect the relative rates of naïve and primed spacer acquisition, which could be mentioned when discussing the results of Figure 1.

We think it is very likely that expression levels of the different *cas* genes impacts the rate of primed or naïve spacer acquisition. Presumably, the reason that other groups have detected naïve spacer acquisition is because they express *cas1* and *cas2* at extremely high levels. We now mention this in the text. Importantly, expression of *cas1* and *cas2* in our experiments is the same in all strains, regardless of whether priming is occurring. Hence, we can directly compare the efficiency of primed and naïve spacer acquisition in these experiments.

6. In the Results section: "Hence, even though we sequenced the first acquired spacer in each array, this spacer may have been acquired as a result of primed adaptation using a newly acquired spacer in the other array." It is unclear from this statement whether the authors observed multiple spacers acquired into an individual CRISPR array. Although a +2 band may not be visible by PCR, reads with a second spacer acquired may be observed in the high-throughput sequencing (depending on how the sequencing was done, see minor concern #1 below). Were +2 products observed in the HTS data? If not, it seems highly unlikely that two acquisition events occurred within two different CRISPRs, but never occurred within an individual CRISPR. Or is the absence of +2 products in the HTS data due to some aspect of sample preparation/sequencing that was not included in the Methods? It would be helpful to describe this result in more detail.

We computationally restricted our analyses to arrays with a single spacer addition. It is clear from the gel of the PCR products (Cooper *et al*., 2018, Figure S1, region ii) that there are many cells with two or even three added spacers in the CRISPR-II array. This supports the idea that chromosomal pre-spacers, as well as pre-spacers from the *cas3*-expressing plasmid, are acquired due to successive rounds of priming. We also note that the strand and AAG PAM biases on the *cas3*-containing plasmid (Figure 2C) strongly support the idea that some of the spacers in the +1 arrays are acquired following an earlier round of priming. By contrast, for experiments where priming happened on the chromosome, there were no detectable +2 expanded CRISPR-II arrays by PCR, suggesting that successive rounds of priming were rare. We have added a gel image to Figure 3. PCR is not the most sensitive assay, and as we discuss below, there is strong evidence for a low level of multiple rounds of priming, but the gel suggests these events were rare. We now discuss these points in the text.

7. Related to the above comment, if +2 products were not observed, it seems unlikely that the spacers matching to the Cas3 plasmid or the chromosome were due to priming from an original spacer acquired from stp. HTS is far more sensitive than the YFP assay, and it is quite possible that these spacers were acquired through naïve adaptation. The location data in Figure 2c-d does suggest that priming occurred, but in that case, it seems like +2 products should have been observed in the HTS data.

See response to (6).

8. Related to the above comment, for the experiment shown in Figure 3, were spacers acquired from regions far away from the protospacers? Analysis of the frequency of spacer acquisition at sites distal (i.e. >200 kb) from the protospacers could provide information on how frequently spacers were acquired through naïve acquisition. Presumably, these spacers would not have been acquired through priming.

We did detect some pre-spacers in other chromosomal locations. We think these were due to successive rounds of priming. These pre-spacers can be classified into two groups. The first group, which include the most abundantly acquired pre-spacers, are those that match sequences from the *cas3*-expressing plasmid. We assume that these were acquired from the plasmid rather than the chromosome. We also assume these sequences were acquired by priming, based on the strong strand bias and AAG PAM bias. Perhaps a spacer was acquired from the rDNA terminator, which is also present on the plasmid, leading to an additional round of priming. The second group of chromosomal pre-spacers, which are much less frequently acquired, are due to second-round priming events either from sequences that match the *cas3*-containing plasmid or repetitive sequences close to the initially targeted protospacer. These pre-spacers have the hallmarks of priming (strand and biases, and decreasing usage frequency as a function of distance from the protospacer). Because there are duplicated sequences scattered around the chromosome, we cannot confidently quantify naïve spacer acquisition by analyzing chromosomal pre-spacers far from the targeted protospacer.

9. For spacers containing an internal AAG sequence, did the authors examine whether the spacer that could originate from that internal AAG was relatively enriched? This type of analysis may indicate whether Cas3 stalls at PAM sites, which could lead to preferential acquisition of spacers from the "first" PAM encountered out of two PAMs that are in

close proximity, similar to the first model that the authors rule out (competition for Cas3 binding between prespacers in close proximity).

We did look at this but didn't include the data. We observe no significant enrichment/de-enrichment for pre-spacers for which the AAG is the PAM, but falls within another pre-spacer with an AAG PAM. We now discuss this briefly in the text.

10. The results shown in Figure 14 are interesting, but are dependent on an exponential decay model that is based on the frequency of spacer acquisition as a function of distance from the protospacer. It is unclear how this model was derived. The Methods says that the data shown in Figure 11a-b were fit to a "best fit line in Excel". What is the best fit line? This should be shown in Figure 11, and the equation used to fit the curve should be specified in the Methods section. To our eye, the data in Figure 11a-b appear that they may fit to a linear model better than to an exponential decay.

We now include the $R^2$ values (0.51 and 0.66) for the best-fit lines in the text. Note that the *y*-axes in Figure 11+B are on a log scale, so while the data fall on a straight line, they fit an exponential decay model.

Minor concerns:

1. Please provide more information on the length of amplicons sequenced by MiSeq and the number of cycles used for the sequencing. Were all possible amplicon lengths accounted for in the MiSeq data (i.e. could the MiSeq cover both +1 and +2 products)?

We now include these numbers in the methods section. The sequencing reads were long enough to detect both +1 and +2 products.

2. It would be helpful if a simple graph showing the relative number of reads derived from each source were included (e.g. a pie chart divided between stp, cas3 plasmid and chromosome).

Because sequences are shared between the different DNA elements, such a pie-chart would be misleading.

3. In Fig. 2D it is very difficult to see the gray bar showing the region of overlap between the stp and the chromosome. Consider changing the color of this bar to something more distinct from the data points.

We have added changed the bars in panels C and D to orange to make them clearer.

4. In the introduction: "A key feature of all CRISPR-Cas systems is their ability to acquire new immunity elements, in a process known as adaptation" The term "immunity elements" is unnecessarily vague and could be mistaken for other immunity elements, such as *cas* genes. Consider changing to "spacers".

Fixed as suggested.

5. The last two sections of the Methods appear to be duplicated. Although they contain some differences, they describe the same analysis.

The key difference is that we excluded AAG-containing pre-spacers for the second analysis. We have rewritten the second section to highlight the similarities and differences to the first section.

Comments of references:

1. In the introduction: “Targeting of a protospacer by Cascade with a partially or fully complementary spacer leads to recruitment of Cas3, which is both a nuclease and a helicase.” References should be added to this sentence.

We have added relevant references.

2. The authors might consider incorporating a recent study into their discussion of correlation between spacer length and PAM slipping shown in Fig. 9 (PMID 32076262). This study established that Cas1 protects the PAM-end of the prespacer from degradation by 3’-5’ exonucleases, resulting in an asymmetrically processed prespacer that favors integration of the non-PAM end of the prespacer at the leader-repeat junction. Kim et al propose a model in which PAM processing is delayed until after the non-PAM end of the prespacer is integrated, which defines the orientation of integration. In the current study, the authors observed that 32 bp spacers correlate with +1 slipping while 34 bp spacer correlate with -1 slipping. Based on the model from Kim et al., following processing and integration of the non-PAM end, over-processing or incomplete processing of the PAM end leads to 32 bp +1-slipped or 34 bp -1-slipped spacers, respectively. This model could be discussed in the section describing the data in Figure 9.

We have expanded the introduction to include a description of this work as well as the related papers from the Siksnys and Bailey groups. We also now briefly discuss the potential role of imprecise exonucleolytic processing of pre-spacer DNA as a cause for slipping.

3. In several places throughout the text, the authors invoke a model in which Cas3 travels along the DNA following recruitment to Cascade. There is substantial evidence, including in the two papers referenced (Dillard et al. and Redding et al.), that in addition to translocation “along” the DNA, Cas3 can also remain bound to Cascade and instead reel the DNA toward the interference complex. Another paper that directly shows this is PMID 29706536. Based on these observations, the authors may consider removing or rewording the following statements:

- “translocation of Cas3 away from the protospacer-bound Cascade” from the Introduction. This would be a good place to introduce the two different models for Cas3 translocation, either movement along the DNA and away from Cascade or reeling of the DNA toward the Cascade-Cas3 complex.

We have added text to introduce this idea. As described in the model at the end of the paper, we think that the reduction in pre-spacer usage very close to protospacers is connected to a reeling mode of Cas3 action.

- “presumably, Cas3 translocates many times around the circular plasmid.” It is possible that Cas3 may dissociate from Cascade and translocate on its own, or that Cas3 may remain bound to Cascade and reel the entire length of the plasmid. This statement does not account for the latter possibility. It is also unclear whether Cas3 would be able to translocate all the way around the plasmid if Cascade remains bound at the protospacer, which may cause a collision and dissociation of Cas3.

In our model, Cas3 breaks away from Cascade after reeling in ~200 bp of DNA. This would be consistent with the plasmid priming data.

- “presumably reflects a bias in the direction of Cas3 translocation away”. The word “away” could be removed. References could also be added to this sentence, as the concept of directional translocation of Cas3 is well established in the literature.

Fixed, as suggested.

- “It is possible that Cas8e translocates with Cas3 and facilitates nicking at AAG sequences.” This section would be a good opportunity to also discuss the idea that Cas3 may remain bound to Cas8e within the Cascade complex and reel in DNA instead of translocating away from Cascade with Cas8e in tow.

This does not fit with our model because Cas8e bound to Cascade at the protospacer would presumably bind the PAM of the protospacer.

- “Hence, we propose that Cas3 transitions stochastically between a state where it nicks DNA frequently, generating an extended region of single-stranded DNA, and a state where it nicks DNA infrequently, with the average distance traveled until the transition being 100-200 bp.” This is an interesting idea, and may suggest that translocation becomes more processive the further Cas3 is “away” from Cascade. There is evidence in the DNA curtains papers that Cas3 eventually breaks away from Cascade, so perhaps this transition event is associated with Cas3 dissociation from the R-loop complex (e.g. a transition from “reeling” to “translocation along the DNA”.

This is our preferred model. We have modified the text to highlight this.

- The model presented in the Discussion could be adjusted to account for data in Loeff et al.

See responses to earlier comments.

4. The authors might consider comparing their chromosomal priming results with the results in PMID 30975459. In that study, the authors repurposed Cascade-Cas3 from a type I-E system for genome editing in eukaryotic cells. They observed directional, long-range deletions originating at the protospacer. The lengths of deletions observed in that study are comparable to the distances of primed spacer acquisition observed in the current study. Combined, these studies highlight the ability of Cas3 to translocate long stretches (~100 kb) of DNA.

This is a great suggestion. There is now another paper that describes genome editing in human cells with the *E. coli* type I-E system, reaching similar conclusions. We have added text to discuss these two papers.

5. In the results: “In type I-F systems, the only characterized type I systems where Cas3 translocation appears to occur with equal efficiency on both the target and non-target strands, Cas2 is fused to Cas3, such that there are two Cas3 subunits in the Cas1-2-3 complex (Fagerlund et al., 2017).” Please also cite PMID 28438998, which was published back-to-back with Fagerlund et al. This citation also provides evidence for bidirectional degradation of the target DNA in type I-F. As mentioned in the first major concern comment, Shiriaeva et al. 2019 suggested that bidirectional spacer acquisition can also occur from bonafide targets in type I-E (although this has not been observed against plasmid DNA to our knowledge).

Citation added as suggested. While Shiriaeva *et al*. reported a much higher proportion of priming on the target strand then we observed, it still corresponds to a minority of all newly acquired spacers.

6. In the Results: “Cas8e, a subunit of Cascade, binds specifically to AAG PAMs.” This statement could use a reference. Also, Cas8e is rather promiscuous for PAMs, so it is unclear whether Cas8e really helps improve PAM specificity during priming.

We have added references for the AAG preference, and clarified the specificity of Cas8e for AAG and other sequences.

Suggestions on improving readability:

1. The HTML version of the manuscript on the *bioRxiv* website is well laid-out. However, the pdf version of the manuscript was a challenge to read because figures and figure legends were not together. There are a lot of figures in this work, so it would be very helpful if the layout could be updated in a manner where the figures are paired with their respective figure legends.

We have reformatted the paper so that the figures and legends are positioned within the text.

2. Consider combining Figures 12-13, as Figure 12a is required to understand data shown in Figure 13.

We have opted to keep these figures separate since they address different, albeit related points.

**Reviewer #2:**

Stringer et al. present a highly comprehensive study that analyzes the characteristics of primed CRISPR adaptation in the type I-E CRISPR-Cas system of E. coli. The study design is solid and the findings are interesting, deepening our view of primed adaptation and preference of new spacer sequences during priming. The findings are well discussed in view of the large body of available literature on this topic. The study is carried out very thoroughly and with a lot of depth.

Below several suggestion that may improve the paper.

L120 "Based on these factors, we propose a model of primed adaptation in which a Cas3-containing complex travels large distances along the DNA from the protospacer, cutting the DNA at or near every AAG, contributing to the generation of substrates for Cas1-2."

1. I am not sure what is different in this model compared to existing models that have been proposed. The long distances have been determined in a number of studies, for example in studies investigating the Cascade-Cas3 for making genomic deletions (Dolan 2019) and (Dillard 2018), and others. It is fine to say that supporting data for this model has been found. What is wrong with that these days?

It was not our intention to present our model as completely novel. While there are novel aspects, it incorporates data from many other studies. We have modified the text to better reflect this.

2. I am not not sure if the authors mean that the AAG containing strand is actually cleaved, or if it could also be the complement. See below.

This section of the paper does not discuss which strand is cut. We have edited the description of the model to point out that either strand could be cut. If Cas3 makes the cut, it is more likely to be the non-target strand, based on where the majority of Cas3 nicks have been detected close to the protospacer.

L390-L406 suggestion

Perhaps no spacers are acquired from regions close to the protospacer because these regions are first looped out by reeling (Loeff, 2018) before Cas3 breaks off, and starts translocating away from Cascade. This unifying view, for which there is experimental evidence, could be discussed as speculation for the observed effects. The proposed transition between states is fine but very speculative, and should be indicated as such. In L571 looping is again relevant to mention.

This is essentially our model. See our response to comments from Reviewer 1.

L329 In vivo Cas8e also dissociates from Cascade (Vink, 2020)

We now include a mention of this result.

L573-L583

This section is again rather speculative when taken literally. It may be more likely that the effects the authors are seeing is happening on the complement of the AAG PAM. CTT is the sequence that Cas1 recognizes as shown by many. Cas3 has some preference for T-stretches (Kunne 2016). Please be careful in how data from spacer sequencing can be translated to mechanistic insights at the molecular level to prevent causing misunderstandings and misconceptions in the field.

See the response to the earlier comment.

Finally it would be good to include some recent insights that have been obtained on PAM selection of prespacers by the Cas1-Cas2 complex, and the processing of prespacers outside the bound part by host nucleases, as well as spacer integration. This seems not to be discussed in much detail and should be added to improve the manuscript (3-4 references). It is the CTT sequence that is recognized by the Cas1-Cas2 complex, and this becomes relevant when converting and interpreting data from new spacer sequencing efforts into biochemical actions of the individual Cas protein components. The resulting inclusion of the text may make the paper more up to date and accurate.

We now mention these three papers in the introduction (see response to Reviewer 1). We don't include them in the discussion because these processing steps likely occur after the events we are focused on.

**Reviewer #3:**

In the paper **"Characterization of Primed Adaptation in the *Escherichia coli* type I-E CRISPR-Cas System"**, Stringer and colleagues present a study investigating priming by the type I-E CRISPR-Cas system. The authors have established a system that triggers priming along the host genome and were able to measure abundance and distance of each pre-spacer from the protospacer. Usually acquisition on host genomic material leads to autoimmunity and cell death, although the authors do not discuss this. By analysing the pre-spacers, the authors showed first acquisition from priming can be made over distances of >100 kb. They also make some interesting observations, including the location and strand of the pre-spacer, slips and flips of the pre-spacer integration, the effect on pre-spacer usage with an internal PAM, and the sequence importance at the PAM-distal end of the pre-spacer. This is a very well written, solid piece of work. In general, the findings in this paper are in support of previous observations in the type I-E priming and also priming in other systems. Some of the findings, such as the importance of the PAM-distal region and internal PAM, are very interesting observations and further in-depth analysis could provide key mechanistic insight into how pre-spacers are selected. The following are a mix of comments, both major and minor, that I believe would improve the paper, or worth considering in the future.

L62: The author states, "During adaptation, one or more spacer+repeat units are added to the end of a CRISPR array." This suggests new spacers are added to the 3' end of the array. This should be reworded to make clear spacers go at the leader-proximal end.

Fixed as suggested.

L66: At first mention, the Cas1-2 complex should be named in full, Cas1-Cas2.

We changed all instances of Cas1-2 to Cas1-Cas2.

L68: Priming can also occur when the PAM has a mismatch. Some acknowledgement of this should be included.

We now mention this in the introduction.

L95: The authors state that most studies have involved an "optimal PAM". What is an optimal PAM? As opposed to what?

We have modified the text to explain that more than one PAM sequence is typically sufficient to bind Cas proteins, but that there is an optimal sequence (or sequences) that have the highest affinity for Cas proteins.

L104: Brief discussion on how spacer polarity occurs, and therefore prevention of flipping, would have been useful.

See response to Reviewer 1.

L142 and L145: Fluorescence was detected, not adaptation. Result is indicative of adaptation. A direct test, such as PCR amplification of array, would support this claim.

We have modified the text to indicate that we infer that adaptation occurred. We did not attempt to PCR from sorted cells because we had a technical problem collecting the cells. Nonetheless, the assay is well established,

L159: I think it is worthwhile noting here that it is only the first acquired spacer that is analysed.

We have modified the text to indicate this.

L183, L189: Chromosomally acquired pre-spacers should be lethal to the host and also should clear any target plasmid. Some brief discussion on autoimmunity, and also priming adaptation vs interference would be useful.

In these experiments we expect both priming and interference. We do not grow the cells for long after inducing priming/interference, so we presume they are dying. Had we not observed chromosomal priming in the plasmid-based assay, we would have assumed that autoimmunity would prevent us from being able to initiate priming on the chromosome. See also our response to Reviewer 1.

L221: The authors state that adaptation from a plasmid does not appear to decrease as a function of distance from the protospacer. This has been clearly shown in the I-F system (Richter et al 2014).

We have modified the text to clarify that are referring specifically to type I-E systems.

L244: The authors suggest that off-target Cascade binding events do not lead to primed adaptation. A site with a mismatched PAM would be “off-target”, but these can lead to priming.

We would not consider such a site to be “off-target” since this level of sequence match would never occur by chance. We have clarified this in the text.

L249 and elsewhere: Spacers are sequenced from genomic/plasmid DNA that is double-stranded. Spacers should be referred to as bp and not nt.

Fixed as suggested.

L265: This next part is difficult to follow. The pre-spacers obtained from the “pink-box” region is compared between the mhpT- and lacZ+ datasets. Figure 6B compares mhpT- target strand vs lacZ+ non-target strand and shows a similar abundance of specific spacers between the two sets, and the authors comment on the very similar pre-spacer location and abundance (comparing for both the minus strand, right of the protospacer). However, in fig 3 these regions do not appear to have the same abundance, although there is no scale to properly compare the graphs. Additionally, the mhpT- pre-spacers must have come from flipping to be on the target upstream strand, which may affect the pre-spacer selection criteria and therefore may not be comparable to the lacz+ dataset.

We are comparing pre-spacer usage on the target and non-target strands for regions upstream of the protospacer. While the absolute level of pre-spacer usage is much higher for the non-target strand, the relative usage frequency for different pre-spacers is the same whether they are on the target or non-target strand. This is not flipping because we are looking upstream of the protospacer. We have clarified this in the text.

L306 (also L353): Based on the proposed hierarchy of selectivity within the PAM and fig 7C, do the authors have some comment over why “-2” slips had low frequency and “+2” slips had increased frequency? Perhaps there is some conservation of bases in the “+2” slips in their second and third positions? This may partly address whether slipping occurs by chance/error, or if there are some underlying sequence preferences that drive this process.

This is an interesting point that we hadn't considered. +2 slips are enriched for GNG PAMs, consistent with the idea that +2 may be preferable to -2 because it allows for a G in the third position of the PAM. However, you would also expect to see the same thing happening with +1 and -1 slips, but there is no enrichment for G in the third position of the PAM for +1 slips. Given that the data are conflicting, we have opted to add text to suggest that sequence preferences may be a factor in determining the relative frequencies of different types of slip, as may structural features of the enzyme that processes the PAM during priming.

L318: It might worth mentioning that the adaptation complex (Cas1-Cas2, Cas1—Cas2-3) specifically detects the PAM in the Cas1 subunit (Wang et al. 2016, Nunez et al. 2016), and is involved in specific cleavage at PAMs and integration (Wang et al. 2016; Fagerlund et al. 2017).

We now mention in the introduction that Cas1-Cas2 recognizes the PAM. We have not discussed Cas1 cleavage at the PAM because there are conflicting data on this activity for *E. coli* Cas1 (e.g. Ramachandran *et al*., 2020; Kim *et al*., 2020).

L355: The authors need to be clearer here. When they state, "pre-spacer usage for the canonical pre-spacers", are they explicitly meaning only the flipped spacers with no slip? I don't think canonical is the best term to refer to these (L353) as it confuses with the majority of pre-spacers that aren't slipped and flipped.

The observation is that flipping occurs more frequently with slipped pre-spacers than it does with non-slipped pre-spacers. This is presumably because the PAM is already mismatched in a slipped pre-spacer, so the penalty for flipping is much lower than if the pre-spacer had an AAG PAM. We have reworded the text to clarify this.

L357: The authors state "pre-spacers generated by slipping events are more likely to be flipped because they have a suboptimal PAM". Where is the data that supports this?

This is now more clearly framed as speculation.

L394: The authors suggest pre-spacer usage frequency is an inherent property of the pre-spacer and not its location. However, the authors previously show that usage is a function of distance from the target. How are these observations consistent? Fig 10 A blue box clearly shows acquisitions are greater in lacZ+ than coda+.

The observation here is that pre-spacer usage is not related to *protospacer* location. Distance from the protospacer is hugely important, but the identity of the protospacer is not. We have changed "location" to "identity" to clarify this point.

L425: This is a really nice discovery. I am interested by the frequency of the internal spacer usage compared to what was expected. Was this also lower?

See response to Reviewer 1, who wondered if it was higher!

L428: Remove "that includes a site for easy introduction of potential pre-spacers flanked by an AAG PAM". Plasmid stp2 already has the pre-spacer, it is not the original plasmid spacers are cloned into.

Done.

L431: Why was position 20 altered, particularly when the earlier positions were later shown to be more important? Also mentioned in the L506 comment, it would have been interesting to see the effects of moving the PAM along the pre-spacer.

We agree it would have been better to look earlier in the sequence. In part, we wanted to include enough upstream sequence that we could test the dependence on melting temperature.

L441: What about the frequency level of the new pre-spacer created with the internal AAG?

We did not look at this because there is nothing to compare it to.

L485: This is an interesting proposal. If Cas3 does nick all PAMs it encounters, how does Cas3 single-stranded DNA helicase activity fits in with this model. Are all nicked single strands then rehybridized with the opposite strand? Perhaps very short fragments that occur between PAMs aren't well rehybridized and making the pre-spacer a poor substrate for integration?

We also thought that short fragments between PAMs might not rehybridize. However, data in Figure 13 argue against this.

L506: I'm not convinced this is the best assay to investigate this hypothesis. Why not move the PAM along the pre-spacer while making minimal base changes? Comparing mut3 to mut2 may give some further insight?

mut3 and mut2 behave similarly. Moving the PAM along the pre-spacer does not allow us to distinguish between a distance-dependent effect and an effect of melting temperature.

L539: The authors observation of base preference at the 3' end of the protospacer is interesting. By changing the two bases distal to the PAM the pre-spacer became notably under-used in acquisition. It would be interesting to see comparison of mut5 to mut4 as there are fewer differences between these pre-spacers. I wonder if this is related to affinity to the Cas1 DNA binding pocket? The hypothesis would have been supported further if the pre-spacer was mutated to bases that were enriched in fig. 14A, predicting an increased frequency of the pre-spacer being acquired.

The relative efficiency of acquisition of mut4 and mut5 can be inferred from Figure 14. mut4 has no detectable difference compared to the unmodified pre-spacer, whereas mut5 is acquired much less efficiently. We agree that it would be interesting to test additional 3' end sequences to compare to predictions based on the data shown in Figure 14A+B.

L574 and L577: There appears to be a contradiction in Cas3 nicking state. On one line the authors say Cas3 transitions to an infrequent nicking state, but then say that it is likely Cas3 nicks at almost every PAM. Can the authors clarify this?

We consider nicking at every PAM to be infrequent, as compared to nicking indiscriminately.

L672: The serendipitously priming function of these plasmids is an unusual set up. I'm not sure what the authors mean that it has a spacer, but not a canonical spacer. The authors propose the array is transcribed in the reverse direction. However, wouldn't this make the down-stream repeat only 8 bp and missing the stem-loop handle for Cas6 binding and processing, which is required for Cascade formation? This is also another case where interference should destroy the plasmid with the array and therefore stop any CRISPR response.

These plasmids were made incorrectly. We had intended to introduce a self-targeting spacer between the two repeat sequences (note that the upstream repeat sequence only includes the 5' handle). However, we used the plasmids before introducing the self-targeting spacer. Surprisingly, they demonstrated robust priming, so we opted to use these data rather than redoing the experiments after cloning the self-targeting spacer. The plasmid has a full repeat sequence that is intended to be downstream of the introduced spacer (the spacer we didn't add!). We think this full repeat serves as an upstream repeat for a sequence immediately downstream that acts as a spacer, even though it is not flanked by a downstream repeat. We think the transcription terminator and associated hairpin act in place of the usual 3' handle. Fortuitously, the pseudo-spacer sequence is duplicated in the plasmid, with the putative protospacer having an AGT PAM.

L923: Authors should indicate whether oligos are 5' – 3'.

Done.

L935: What do the error bars represent? What is *n*?

Fixed, and fixed for Figures 12 and 14.

L956: There are no values on the graph.

This refers to the values on the *y*-axis. We didn't include actual numbers because they are meaningless; only the relative numbers matter.

Fig 2: Shouldn't panel A have grey boxes similar to C? Is there uncertainty where pre-spacers were derived from in this plasmid as well?

Since priming was initiated on this plasmid, we assume that most of the pre-spacers that match other sequences (i.e. the *cas3*-expressing plasmid or the chromosome) were derived from the stp.

Fig 6: I suggest flipping the lacZ+ diagram so each quadrant is similar to the graph below. i.e. Top left for both diagram and graph is non-target strand, upstream. I think this is easier to understand and the fact that lacz+ and mhpT- target are on different strands is irrelevant.

We have opted to keep the orientation as it is. This highlights the fact that the two protospacers are on opposite strands. It is unlikely that this is relevant for the data in Figure 6, but it might be, e.g. if orientation relative to the replication fork impacts primed adaptation.

Fig 10: Panel A, it would be useful to show where the protospacer is.

Done.
