## Supplementary material for "Characterization of Primed Adaptation in the *Escherichia coli* type I-E CRISPR-Cas System": Table S1

**Table S1. Oligonucleotides and synthesized dsDNA used in this study.**

| Name | Sequence (5′ to 3′) |
|---|---|
| JW472 | CCGACGCGCAGTTTA |
| JW473 | CACGTTGTGTTTTCATGC |
| JW6518 | CAGCGGGGATAAACC |
| JW7816 | TCGTCGGCAGCGTCAGATGTGTATAAGAGACAGAAGGTTGGTGGGTTGTTTTTATGGG |
| JW7817 | GTCTCGTGGGCTCGGAGATGTGTATAAGAGACAGTTATCAATTACAACCGACAGGGAGCC |
| JW7818 | TCGTCGGCAGCGTCAGATGTGTATAAGAGACAGAAAGTTGGTAGATTGTGACTGGC |
| JW7819 | GTCTCGTGGGCTCGGAGATGTGTATAAGAGACAGCAACAGCAGCACCCATGAC |
| JW7820 | AATGATACGGCGACCACCGAGATCTACACGCGTAAGATCGTCGGCAGCGTC |
| JW8010 | GCGCGGGGAACTCGAGCAGTCGCGCTTTGTCGAAACCGTTGCTGCCGGTTTATCCCCGC |
| JW8011 | GCGCGGGGAACTCGAGGTGCCAGGGCATACAAAACGCTTTGCCCACGGTTTATCCCCGC |
| JW8053 | AATGATACGGCGACCACCGAGATCTACACTCCAGGTATCGTCGGCAGCGTCAGATGTG |
| JW8054 | CAAGCAGAAGACGGCATACGAGATGGATTCACGTCTCGTGGGCTCGGAGATGTG |
| JW8057 | CAAGCAGAAGACGGCATACGAGATCGCATTAGGTCTCGTGGGCTCGGAGATGTG |
| JW8062 | CAAGCAGAAGACGGCATACGAGATATGGACTCGTCTCGTGGGCTCGGAGATGTG |
| JW8476 | CAAGCAGAAGACGGCATACGAGATTAAGGCGAGTCTCGTGGGCTCGGAGATGTG |
| JW8477 | CAAGCAGAAGACGGCATACGAGATCGTACTAGGTCTCGTGGGCTCGGAGATGTG |
| JW8478 | CAAGCAGAAGACGGCATACGAGATAGGCAGAAGTCTCGTGGGCTCGGAGATGTG |
| JW8479 | CAAGCAGAAGACGGCATACGAGATTCCTGAGCGTCTCGTGGGCTCGGAGATGTG |
| JW8480 | CAAGCAGAAGACGGCATACGAGATGGACTCCTGTCTCGTGGGCTCGGAGATGTG |
| JW8481 | CAAGCAGAAGACGGCATACGAGATTAGGCATGGTCTCGTGGGCTCGGAGATGTG |
| JW8485 | CAAGCAGAAGACGGCATACGAGATCGAGGCTGGTCTCGTGGGCTCGGAGATGTG |
| JW9009 | ATTGGGCCAGCTAAATCG |
| JW9010 | GGCTCATTATACCAGTCAGGACGTTGGGAAGAGGCCGCTCAAACAGGTAAAAAAGACACC |
| JW9016 | GCTAAATCGATGGGATGTGGCTTGCTATCTTTGGCTCCACTGTGATAGACAGCTGCATGCAT |
| JW9017 | AGAACTGGCTCATTATACCAGTCAGGACGTTGGGAAGAGGCCGCGTGTAGGCTGGAGCTG |
| JW9066 | TGTGGAATTGTGAGCGGATAACAATTTCACACAGGAAACAGCTTAGACAGCTGCATGCAT |
| JW9067 | TTCCTTACGCGAAATACGGGCAGACATGGCCTGCCCGGTTATTAGTGTAGGCTGGAGCTG |
| JW9076 (dsDNA) | TGTTTGACAGCTTATCATCGATTTGACAGCTAGCTCAGTCCTAGGTATAATGCTAGCATAAACCGCGGTACTTAGCTCCTCAGATTAGGATTGCGGAGAATAACA |

| | |
|---|---|
| | ACCGCCGTTCTCATCGAGTAATCTCCGGATATCGACCCATAACGGGCAATGATAAAAGGAGTAACCTGTGAAAAAGATGCAATCTATCGTACTCGCACTTTCCCTGGTTCTGGTCGCTCCCATGGCAGCACAGGCTGCGGAAATTACGTTAGTCCCGTCAGTAAAATTACAGATAGGGAGGAGCGATCCTGCAGTGTTCCCCGCGCCAGCGGGGATAAACCGAGGGAACTGCCAGGCATCAAATAAAACGAAAGGCTCAGTCGAAAGACTGGGCCTTTCGTTTTATCTGTTGTTTGTCGGTGAACGCTCTCCTGAGTAGGACAAGCTTGGCTGTTTTGGCGGA |
| JW10009 | TGAGTAGGACAAGCTTACTAATTATAATAGAAGCCAGATACTAAAAAGCTTGGCTGTTTT |
| JW10010 | AAAACAGCCAAGCTTTTTAGTATCTGGCTTCTATTATAATTAGTAAGCTTGTCCTACTCA |
| JW10011 | TGAGTAGGACAAGCTTACTAATTATAATAGAAACCAGATACTAAAAAGCTTGGCTGTTTT |
| JW10012 | AAAACAGCCAAGCTTTTTAGTATCTGGTTTCTATTATAATTAGTAAGCTTGTCCTACTCA |
| JW10013 | TGAGTAGGACAAGCTTGCCGGCTGCAGCAGAAGCCAGATACTAAAAAGCTTGGCTGTTTT |
| JW10014 | AAAACAGCCAAGCTTTTTAGTATCTGGCTTCTATTATATATAGTAAGCTTGTCCTACTCA |
| JW10015 | TGAGTAGGACAAGCTTGCCGGCTGCAGCAGAAACCAGATACTAAAAAGCTTGGCTGTTTT |
| JW10016 | AAAACAGCCAAGCTTTTTAGTATCTGGCTTCTATTATATATAGTAAGCTTGTCCTACTCA |
| JW10017 | TGAGTAGGACAAGCTTACTAATTATAATAGAAACCAGATACCCCGAAGCTTGGCTGTTTT |
| JW10018 | AAAACAGCCAAGCTTCGGGGTATCTGGTTTCTATTATAATTAGTAAGCTTGTCCTACTCA |
| JW10021 | TGAGTAGGACAAGCTTACTAATTATAATAGAAACCAGATACCCAAAAGCTTGGCTGTTTT |
| JW10022 | AAAACAGCCAAGCTTTTGGGTATCTGGTTTCTATTATAATTAGTAAGCTTGTCCTACTCA |
