## Supplementary material for "Characterization of Primed Adaptation in the *Escherichia coli* type I-E CRISPR-Cas System": Marked-up copy showing changes to the original submission in response to the reviews

Intriguingly, many pre-spacers did not map to the stp, suggesting another source for the acquired spacers. ~~We determined whether any of the unmapped pre-spacers map to the *cas3*-expressing plasmid or to the chromosome. Many~~ Most of the pre-spacers that did not map to the stp ~~reads map~~ mapped uniquely to the *cas3*-expressing plasmid (Figure 2C), although it is important to note that much of the sequence of this plasmid is identical to that of the stp, and reads that match shared regions were mapped to the stp. Pre-spacer usage on the *cas3*-expressing plasmid was strongly biased to one strand and to pre-spacers with an AAG PAM. Hence, we speculated that pre-spacers mapping uniquely to the *cas3*-expressing plasmid represent a second round of primed adaptation.

Even though we sequenced the first acquired spacer in each array, this spacer may have been acquired as a result of primed adaptation using a newly acquired spacer in the other array. Consistent with multiple rounds of adaptation occurring in individual cells, we previously showed that primed adaptation under these conditions often leads to successive rounds of expansion of the CRISPR-II array (Cooper et al., 2018).

We also detected a small number of pre-spacers that mapped uniquely to the chromosome. We reasoned that these sequences could have been acquired due to multiple rounds of primed adaptation if the first acquired spacer matched a sequence shared between the stp and the chromosome. ~~Unexpectedly, a small proportion of the reads mapped uniquely to the chromosome. We speculated that pre-spacers mapping uniquely to the *cas3*-expressing plasmid or the chromosome represent a second round of primed adaptation. This is possible because the strain we used contains both the CRISPR-I and CRISPR-II arrays. Hence, even though we sequenced the first acquired spacer in each array, this spacer may have been acquired as a result of primed adaptation using a newly acquired spacer in the other array.~~ Consistent with this hypothesis, pre-spacers mapping uniquely to the chromosome were found adjacent to sequences shared with the stp, i.e. the *araC* gene (Figure 2D), or the transcription terminators of rRNA loci. Thus, our data indicate that primed adaptation from chromosomal pre-spacers can be readily detected using this approach.

AAG-associated pre-spacers account for 87.7% (lacZ+) and 95.7% (*mhpT*-) of all 32 ~~nt~~ bp pre-spacer usage. Similarly, the majority of 34 ~~nt~~ bp pre-spacer usage was associated with AAG PAMs; hence, these pre-spacers are equivalent to efficiently used 33 ~~nt~~ bp pre-spacers, but with an additional base at the PAM-distal end. These AAG-associated pre-spacers account for 79.7% (*lacZ*+) and 86.7% (*mhpT*-) of all 34 ~~nt~~ bp pre-spacer usage. There is also a significant enrichment for 32 ~~nt~~ bp pre-spacer usage associated with +1 slips, and for 34 ~~nt~~ bp pre-spacer usage associated with -1 slips (Figure 9), indicating that slipping sometimes involves maintaining the PAM-distal boundary while either shortening or lengthening the spacer. The equivalent observation has been made for type I-B, type I-C and type I-F CRISPR-Cas systems (Li et al., 2017; Rao et al., 2017; Staals et al., 2016).

Raw Illumina sequencing data are available from EBI ArrayExpress using accession number E-MTAB-8700.

This work was supported by National Institutes of Health grants R01GM122836 (JTW) and R21AI126416 (JTW).

**Table 1. Strains used in this study.**

| **Name** | **Description** | **Source** |
|---|---|---|
| MG1655 | Wild-type *E. coli* | (Blattner et al., 1997) |
| CB386 | MG1655 [Δ*cas3* P*cse1*]::[*cat* PJ23119] | (Luo et al., 2014) |
| AMD536 | MG1655 [Δ*cas3* P*cse1*]::[ PJ23119] | (Cooper et al., 2018) |
| AMD688 | MLS1003 [Δ*cas3* P*cse1*]::[ PJ23119] | (Cooper et al., 2018) |
| 1XDNAi | MG1655 Δ*araC-araBAD*, ΔP*lacI*:*lacI*, Δ*cas*-CRISPR::pACT-01 ($kan^-$) | (Caliando and Voigt, 2015) |
| AMD671 | MG1655 Δ*araC-araBAD*, ΔP*lacI*:*lacI*, Δcas-CRISPR::pACT-01, $cas1^+$ $cas2^+$, Δ*lacZ*::*thyA* | This Study |
| AMD052 | MG1655 Δ*thyA* | (Stringer et al., 2012) |

**Table 2. Plasmids used in this study.**

| Name | Description | Source |
|---|---|---|
| pBAD24 amp | Empty pBAD24 amp | (Guzman et al., 1995) |
| pBAD33 cam | Empty pBAD33 cam | (Guzman et al., 1995) |
| pAMD179 | Parent vector for cloning crRNAs | (Cooper et al., 2018) |
| pAMD189 | stp; pAMD179 expressing a self-targeting crRNA | (Cooper et al., 2018) |
| pAMD191 | pBAD33-*cas3* | (Cooper et al., 2018) |
| pCB380 | pcrRNA.con-*lacZ* | (Luo et al., 2014) |
| pAMD211 | pAMD179 expressing a crRNA targeting *mhpT* | This Study |
| pAMD212 | pAMD179 expressing a crRNA targeting *codA* | This Study |
| pSDS009 | Parent vector for cloning pre-spacers | This Study |
| pSK013 | pSDS009 stp2-mut1 pre-spacer | This Study |
| pSK014 | stp2; pAMD179 with an added pre-spacer | This Study |
| pSK015 | pSDS009 stp2-mut2 pre-spacer | This Study |
| pSK035 | pSDS009 stp2-mut3 pre-spacer | This Study |
| pSK016 | pSDS009 stp2-mut5 pre-spacer | This Study |
| psK017 | pSDS009 stp2-mut4 pre-spacer | This Study |

**Table 3. Oligonucleotides and synthesized dsDNA used in this study.**

| Name | Sequence (5′ to 3′) |
|---|---|
| JW472 | CCGACGCGCAGTTTA |
| JW473 | CACGTTGTGTTTTCATGC |
| JW6518 | CAGCGGGGATAAACC |
| JW7816 | TCGTCGGCAGCGTCAGATGTGTATAAGAGACAGAAGGTTGGTGGGTTGTTTTTATGGG |
| JW7817 | GTCTCGTGGGCTCGGAGATGTGTATAAGAGACAGTTATCAATTACAACCGACAGGGAGCC |
| JW7818 | TCGTCGGCAGCGTCAGATGTGTATAAGAGACAGAAAGTTGGTAGATTGTGACTGGC |
| JW7819 | GTCTCGTGGGCTCGGAGATGTGTATAAGAGACAGCAACAGCAGCACCCATGAC |
| JW7820 | AATGATACGGCGACCACCGAGATCTACACGCGTAAGATCGTCGGCAGCGTC |
| JW8010 | GCGCGGGGAACTCGAGCAGTCGCGCTTTGTCGAAACCGTTGCTGCCGGTTTATCCCCGC |
| JW8011 | GCGCGGGGAACTCGAGGTGCCAGGGCATACAAAACGCTTTGCCCACGGTTTATCCCCGC |
| JW8053 | AATGATACGGCGACCACCGAGATCTACACTCCAGGTATCGTCGGCAGCGTCAGATGTG |
| JW8054 | CAAGCAGAAGACGGCATACGAGATGGATTCACGTCTCGTGGGCTCGGAGATGTG |
| JW8057 | CAAGCAGAAGACGGCATACGAGATCGCATTAGGTCTCGTGGGCTCGGAGATGTG |
| JW8062 | CAAGCAGAAGACGGCATACGAGATATGGACTCGTCTCGTGGGCTCGGAGATGTG |
| JW8476 | CAAGCAGAAGACGGCATACGAGATTAAGGCGAGTCTCGTGGGCTCGGAGATGTG |
| JW8477 | CAAGCAGAAGACGGCATACGAGATCGTACTAGGTCTCGTGGGCTCGGAGATGTG |
| JW8478 | CAAGCAGAAGACGGCATACGAGATAGGCAGAAGTCTCGTGGGCTCGGAGATGTG |
| JW8479 | CAAGCAGAAGACGGCATACGAGATTCCTGAGCGTCTCGTGGGCTCGGAGATGTG |
| JW8480 | CAAGCAGAAGACGGCATACGAGATGGACTCCTGTCTCGTGGGCTCGGAGATGTG |
| JW8481 | CAAGCAGAAGACGGCATACGAGATTAGGCATGGTCTCGTGGGCTCGGAGATGTG |
| JW8485 | CAAGCAGAAGACGGCATACGAGATCGAGGCTGGTCTCGTGGGCTCGGAGATGTG |
| JW9009 | ATTGGGCCAGCTAAATCG |
| JW9010 | GGCTCATTATACCAGTCAGGACGTTGGGAAGAGGCCGCTCAAACAGGTAAAAAAGACACC |

| | |
|---|---|
| JW9016 | GCTAAATCGATGGGATGTGGCTTGCTATCTTTGGCTCCACTGTGATAG ACAGCTGCATGCAT |
| JW9017 | AGAACTGGCTCATTATACCAGTCAGGACGTTGGGAAGAGGCCGCGTG TAGGCTGGAGCTG |
| JW9066 | TGTGGAATTGTGAGCGGATAACAATTTCACACAGGAAACAGCTTAGA CAGCTGCATGCAT |
| JW9067 | TTCCTTACGCGAAATACGGGCAGACATGGCCTGCCCGGTTATTAGTG TAGGCTGGAGCTG |
| JW9076 (dsDNA) | TGTTTGACAGCTTATCATCGATTTGACAGCTAGCTCAGTCCTAGGTAT AATGCTAGCATAAACCGCGGTACTTAGCTCCTCAGATTAGGATTGCG GAGAATAACAACCGCCGTTCTCATCGAGTAATCTCCGGATATCGACC CATAACGGGCAATGATAAAAGGAGTAACCTGTGAAAAAGATGCAAT CTATCGTACTCGCACTTTCCCTGGTTCTGGTCGCTCCCATGGCAGCAC AGGCTGCGGAAATTACGTTAGTCCCGTCAGTAAAATTACAGATAGGG AGGAGCGATCCTGCAGTGTTCCCCGCGCCAGCGGGGATAAACCGAGG GAACTGCCAGGCATCAAATAAAACGAAAGGCTCAGTCGAAAGACTG GGCCTTTCGTTTTATCTGTTGTTTGTCGGTGAACGCTCTCCTGAGTAG GACAAGCTTGGCTGTTTTGGCGGA |
| JW10009 | TGAGTAGGACAAGCTTACTAATTATAATAGAAGCCAGATACTAAAAA GCTTGGCTGTTTT |
| JW10010 | AAAACAGCCAAGCTTTTTAGTATCTGGCTTCTATTATAATTAGTAAGC TTGTCCTACTCA |
| JW10011 | TGAGTAGGACAAGCTTACTAATTATAATAGAAACCAGATACTAAAAA GCTTGGCTGTTTT |
| JW10012 | AAAACAGCCAAGCTTTTTAGTATCTGGTTTCTATTATAATTAGTAAGC TTGTCCTACTCA |
| JW10013 | TGAGTAGGACAAGCTTGCCGGCTGCAGCAGAAGCCAGATACTAAAA AGCTTGGCTGTTTT |
| JW10014 | AAAACAGCCAAGCTTTTTAGTATCTGGCTTCTATTATATATAGTAAGC TTGTCCTACTCA |
| JW10015 | TGAGTAGGACAAGCTTGCCGGCTGCAGCAGAAACCAGATACTAAAA AGCTTGGCTGTTTT |
| JW10016 | AAAACAGCCAAGCTTTTTAGTATCTGGCTTCTATTATATATAGTAAGC TTGTCCTACTCA |
| JW10017 | TGAGTAGGACAAGCTTACTAATTATAATAGAAACCAGATACCCCGAA GCTTGGCTGTTTT |
| JW10018 | AAAACAGCCAAGCTTCGGGGTATCTGGTTTCTATTATAATTAGTAAG CTTGTCCTACTCA |
| JW10021 | TGAGTAGGACAAGCTTACTAATTATAATAGAAACCAGATACCCAAAA GCTTGGCTGTTTT |
| JW10022 | AAAACAGCCAAGCTTTTGGGTATCTGGTTTCTATTATAATTAGTAAGC TTGTCCTACTCA |

1057

### FIGURE LEGENDS

**Figure 1. Primed adaptation is >1000-fold more efficient than naïve adaptation. (A)** Schematic showing the *yfp* reporter construct of strain AMD688 used to quantify adaptation. The genome of the parent strain has a single CRISPR repeat embedded within a *yfp* open reading frame such that the repeat causes a frame-shift, preventing translation of YFP. Expansion of the CRISPR array by a single repeat and spacer restores the frame, permitting translation of YFP, leading to detectable fluorescence. **(B)** Percentage of $YFP^+$ cells for an *E. coli* strain containing the *yfp* reporter (AMD688), either the stp (pAMD189; "crRNA +") or an equivalent empty vector (pBAD24; "crRNA -"), and either a *cas3*-expressing plasmid (pAMD191; "cas3 +") or an equivalent empty vector (pBAD33; "cas3 -"), following addition of arabinose to the cultures. Values plotted represent the mean of three independent biological replicates. Error bars represent one standard deviation from the mean.
